# A line attractor maintains aggressiveness during feeding in “hangry” mice

**DOI:** 10.1101/2025.10.16.682711

**Authors:** Jineun Kim, Aditya Nair, Nestor Coria, Sophia Huynh, Amit Vinograd, Jingyue Xu, Mengyu Liu, David J. Anderson

**Affiliations:** Division of Biology and Biological Engineering, California Institute of Technology; Tianqiao and Chrissy Chen Institute of Neuroscience, California Institute of Technology; Howard Hughes Medical Institute Pasadena, CA, USA; Lee Kong Chian School of Medicine, Nanyang Technological University, Singapore; Institute of Molecular and Cellular Biology, Agency of Science, Technology and Research, Singapore; Department of Bioengineering, Stanford University, Stanford, CA

**Keywords:** Dynamical systems analysis, interacting internal states, hunger, aggression, emotion, homeostasis, flexible innate behaviors, state-dependent modulation

## Abstract

Aggression evolved to protect resources such as food from competitors, but animals must balance fighting and feeding so that they facilitate rather than hinder re-establishment of energy homeostasis. How this balancing is computed is not well understood. We have approached this problem at the level of neural population-coding by examining the effect of progressive starvation on a hypothalamic line attractor that encodes an internal state of aggressiveness. Moderate fasting yielded “hangry” mice, decreasing attack latency and increasing attack frequency. In parallel, line attractor ramping rate and stability were increased, suggesting that hunger enhances aggressiveness by modifying neural dynamics. In contrast, prolonged starvation inhibited aggression and eliminated the line attractor. In satiated mice, titrated acute chemogenetic activation of arcuate AgRP neurons recapitulated the biphasic effects of progressive starvation, suggesting that a continuous increase in hunger exerts bi-directional influences at different intensities. When confronted with food and an intruder, hangry mice alternated between feeding and fighting. During eating, population neural activity moved out of the line attractor while activity in the attractor dimension remained unchanged. Following feeding, activity rapidly relaxed back into the attractor and aggression resumed. Thus, the line attractor may serve to keep hungry animals primed for aggression during intermittent feeding bouts.

## INTRODUCTION

Aggression evolved to protect resources essential for individual survival and reproduction such as food, territory, mating partners, or offspring^1^. However, fighting is mutually exclusive with behaviors that mediate resource acquisition and consumption. Thus, for example, hungry animals must dynamically adjust their choice between feeding and fending off competitors for food so that fighting facilitates, but does not ultimately interfere with, re-establishment of energy homeostasis^2–5^. This choice likely depends upon multiple factors such as residual energy reserves, quantity and quality of food resources and perceived aggressiveness of the competitor. How this dynamic decision-making is computed by the brain to optimize flexible behavioral prioritization^6^ is not well understood.

As a first step towards addressing this question, we have investigated how aggressive behavior and brain activity are modulated by the duration of starvation. Hunger is well-known to enhance aggressiveness in multiple species^7–10^, and is experienced in humans as “hanger”^11–13^. Surprisingly, this phenomenon has not been convincingly demonstrated in mice. To the contrary, recent data suggest that food deprivation inhibits rather than enhances aggression in mice^5,14^. Older studies have described bi-directional effects of fasting duration on aggression^15^, but characterization of these effects was limited.

Aggression is controlled by estrogen receptor-1 (Esr1)^16^- and progesterone receptor (PR)^17,18^-expressing neurons located in the ventrolateral subdivision of the ventromedial hypothalamic nucleus (VMHvl^19^; reviewed in refs^20–22^). VMH is also a well-known neural hub for controlling feeding and metabolic regulation^23–25^. Whether and how hunger controls aggression via influences on VMHvl^Esr1/PR^ neurons is not clear. A suppressive effect of starvation on aggression was shown to be mediated by VMHvl-projecting BNST neurons that receive inhibitory input from NPY-responsive medial amygdala (MeA) neurons^14^, but the role of VMHvl^Esr1/PR^ neurons was not investigated.

Given the complexity of VMHvl cell types^26–29^, circuitry^25,30–32^ and single-unit activity^19,33,34^, we have asked whether aggressiveness is modulated by food deprivation at the level of population coding and manifolds in neural state-space^35–39^. Attractors are a well-established population coding mechanism for computing cognitive variables^39–44^. Recently, we identified an approximate line attractor^45^ that encodes a scalable and persistent internal state of aggressiveness in VMHvl^46–48^. Other data suggest that attractor dynamics may encode physiological internal states as well^49^. For example, competition between feeding and drinking is reflected in dynamical interactions between apparent basins of attraction for hunger vs. thirst^50,51^.

Here we report that different durations of starvation exert strongly correlated bi-phasic influences on aggressiveness and on VMHvl^Esr1^ line attractor dynamics, an effect recapitulated by titrated short-term chemogenetic activation of arcuate AgRP-expressing “hunger”-promoting neurons^52–54^ in satiated mice. Furthermore, activity in this line attractor remains persistently elevated when “hangry” mice alternate between feeding and attack bouts in a behavioral choice paradigm. These results suggest that modifications of VMHvl line attractor dynamics may underlie the neural coding of “hanger,” and that this attractor may play an adaptive role to keep hungry animals “primed” for attack while replenishing their energy reserves.

## RESULTS

### Metabolic states modulate male aggression in a biphasic manner, establishing hunger-enhanced aggression in mice

To determine the features of aggressive behaviors in male mice that are modulated by metabolic states, we employed the standard resident-intruder (RI) assay^33,55^ and perturbed the resident’s energy states by limiting food availability with different durations of food deprivation before introducing a rival (*ad libitum*-fed) intruder (Fig. 1a). The same resident mice under *ad libitum* feeding conditions served as within-subject controls, using a cross-over design (Fig. 1b). The RI assay itself was performed in the resident’s home cage in the absence of food.

**Fig. 1.**
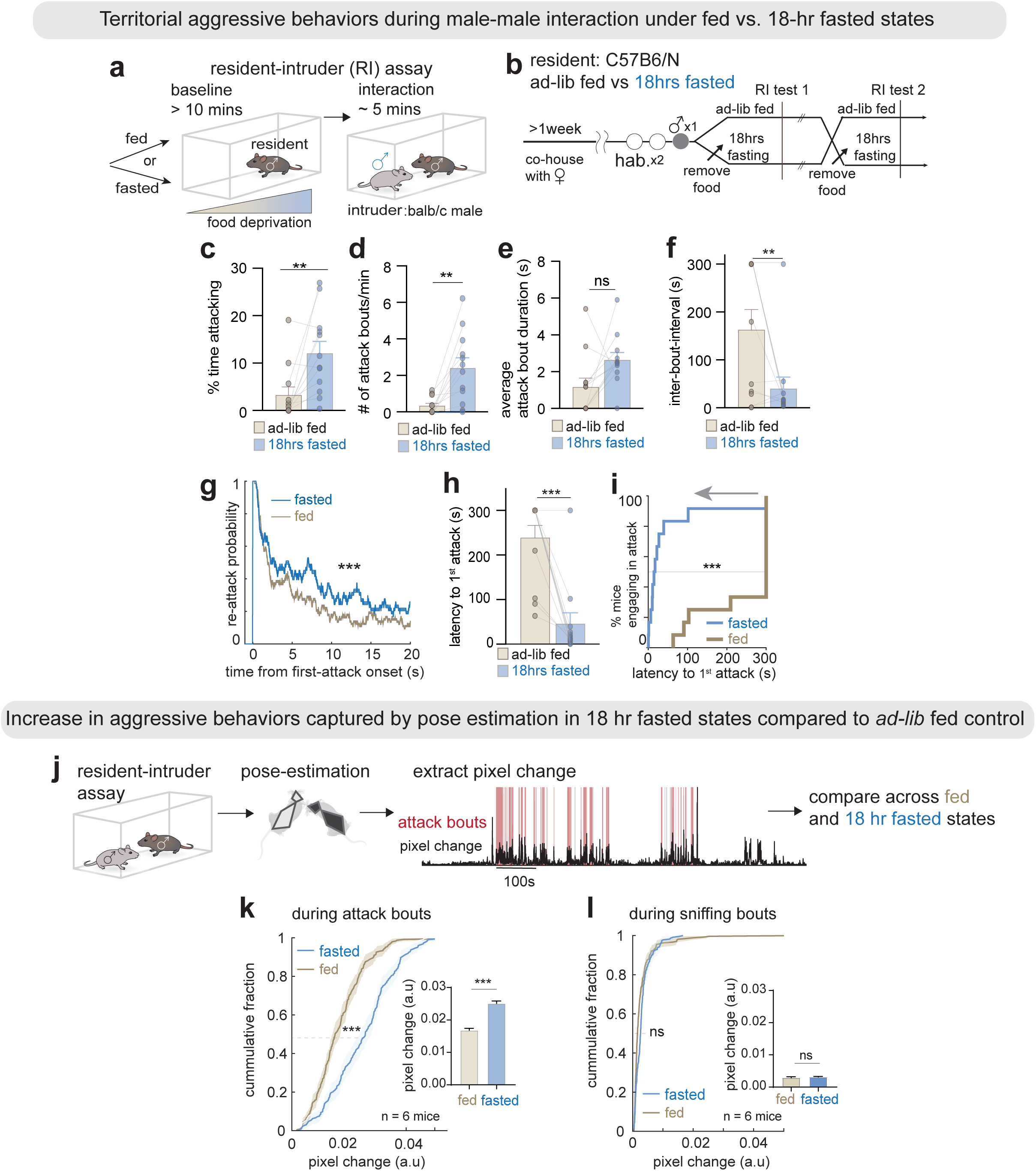
Moderate food deprivation enhances aggression in male mice. (**a**) Schematic of resident-intruder (RI) assay for male-male interactions following *ad libitum*-fed or 18-hr fasting of the resident. (**b**) Schematic of behavioral paradigm to compare aggressive behaviors across different energy states. The order of fed vs. fasted states was counter-balanced across cohorts, with a 2-day interval between conditions (Methods). (**c-f, h**) Quantification of aggression parameters: (**c**) time spent in attack (%); (**d**) number of attack bouts/min; (**e**) average attack duration; (**f**) inter-bout-interval; and (**h**) latency to the first attack. (**g**) Cumulative probability of re-attack after the onset of the first attack bout, across animals (n = 12 mice). (**i**) Empirical cumulative distribution function (ECDF) plot showing latency to the first attack across the *ad libitum*-fed (brown) vs 18-hr fasted state (blue). Data scaled to 100%. (**j**) Schematic illustrating unsupervised analysis of pose estimation features (Methods). The pixel change value associated with the resident mouse during each behavioral bout was computed for each mouse and compared across fed vs 18-hr fasted conditions. (**k-l**) ECDF plots for pixel changes across fed vs. 18-hr fasted states during attack bouts (**k**), and during sniffing bouts (**l**). Bar graphs quantifying the mean pixel change across energy state from 6 mice (right). \*\**p*<0.01, ***p<0.001; ns, not significant. For all figure panels, data are mean ± SEM (n=6 mice in this and all figures unless otherwise indicated). See also **Extended Data Fig. 1**.

Mice that were food-deprived for 18 hours exhibited heightened aggression compared to those fed *ad libitum*, as evidenced by a significant increase in time spent attacking^6^, the number of attack bouts, and a reduction in inter-attack bout interval duration (although there was no change in average attack-bout duration; Fig. 1c-f). In addition, the probability of attack following the initial aggressive encounter increased under starvation (Fig. 1g). Notably, 18h-fasted mice showed a significantly shorter latency to initiate the first attack than the same mice under *ad libitum-*fed conditions (Fig. 1h, i).

We also investigated whether there were additional changes in the kinematics^56^ of aggressive behavior under starvation. We employed machine learning (ML) to quantify the intensity of aggression based on the values of different pose estimation^57^-derived features, using seven key points assigned to each mouse (Fig. 1j; see Methods). Among these features, we found that pixel change, a measure of movement velocity across frame^57^ (Fig. 1j, right-black), was significantly higher during aggression performed by 18-hr-fasted mice compared to *ad libitum*-fed controls (Fig.1k, bar graph). This increase was observed specifically during video frames annotated as attack bouts, but not during sniffing bouts (Fig. 1k vs. 1l). This result corroborates our subjective impression that aggression in mice fasted for 18 hours was more vigorous compared to that in fed animals.

We next sought to reconcile this observation with prior studies demonstrating that food deprivation suppressed aggression in male mice^6,14^. This apparent discrepancy could reflect methodological differences or dynamic biological interactions between hunger and aggression states. To investigate whether the effect of hunger to promote aggressiveness was a linear function of starvation time, we measured aggression after shorter (∼8 hr) or more extended (40 hr) periods of food deprivation. (The durations were chosen based on body weight changes and compatibility with similar zenithal times for performing behavioral assays; Extended Data Fig. 1a, Methods.) Mild starvation (8 hr) did not significantly alter aggressive behavior compared to the fed states (Extended Data Fig. 1b, b). In contrast, prolonged starvation (40 hr) dramatically attenuated aggressive behavior, consistent with previous reports^14,15^, and significantly increased attack latency compared to *ad libitum*-fed mice (Extended Data Fig. 1d, d). Notably, the time spent investigating male intruders was not significantly different across different energy states, although there was a trend to increased sniffing after 40 hours of food deprivation (Extended Data Fig. 1f; see also Fig. 3c and Extended Data Fig. 2a). Thus, food deprivation had a biphasic effect on aggression in mice, with intermediate starvation enhancing and prolonged starvation decreasing aggressive behavior^15^.

We also observed a profound change in male sexual behavior during interactions with female mice under intermediate fasting conditions^6^ (Extended Data Fig. 1g). In the absence of food, male mice exhibited a marked decrease in sexual behavior (such as mounting and intromission toward female intruders) following 18 hours of fasting, suggesting that a deviation from metabolic homeostasis can suppress sexual drive under some conditions (Extended Data Fig. 1h-m). Remarkably, under these conditions, 20% of males even displayed aggressive behavior toward female intruders, a behavior rarely observed under *ad libitum*-fed conditions (Extended Data Fig. 1h-pie chart). This observation underscores the potency of 18 hours of fasting to alter the balance of social behaviors, increasing aggression while decreasing mating.

### Intermediate starvation alters the ramping and stability of line attractor dynamics representing the aggressive state

We next sought to characterize neural correlates underlying the hunger-induced enhancement of aggressiveness; in other words, how is the internal state colloquially termed ‘hanger’ encoded in the brain^46,47^? To this end, we measured the activity of individual VMHvl^Esr1^ neurons expressing GCaMP8m by performing *in vivo* microendoscopic calcium imaging^58^ in mice fed *ad libitum,* and in the same mice following 18 hours of food deprivation. This longitudinal imaging allowed us to track the same population of neurons over several days (see Methods) to examine changes in their relative activity across distinct energy states (Fig. 2a-d). We confirmed that mice with GRIN lens implants and head-mounted miniature microscopes recapitulated the phenotypic readouts of hunger-induced changes in aggressive behaviors (Extended Data Fig. 2a-e).

**Fig. 2.**
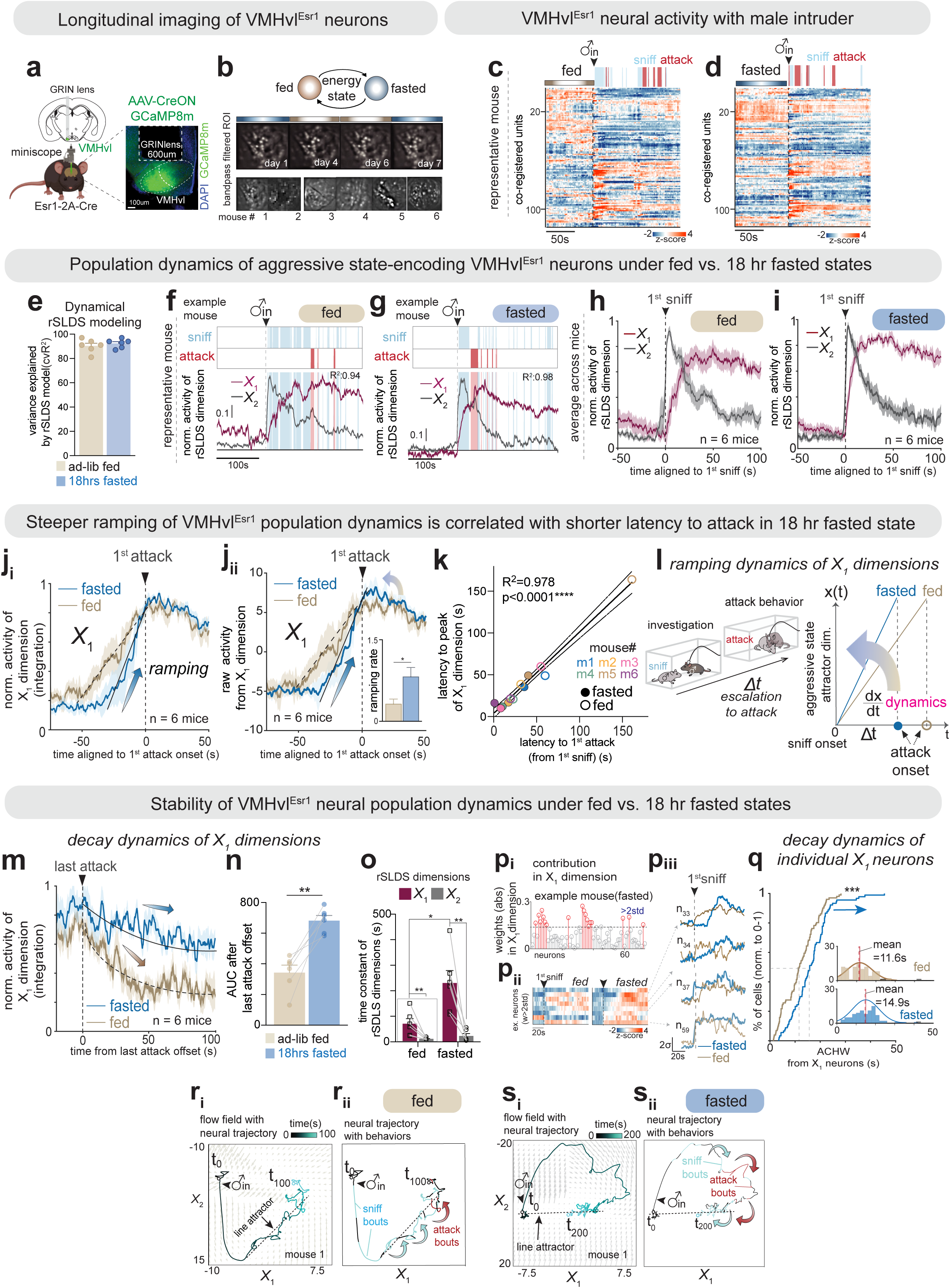
Moderate fasting enhances ramping and persistence in a VMHvl^Esr1^ latent slow mode. (**a**) (left) Imaging preparation and (right) representative histological section showing GCaMP8m expression (green) and GRIN lens implantation site. (**b**) Representative imaging planes from longitudinal *in vivo* calcium imaging; color code indicates energy state (above) and mouse number for analysis (below). (**c-d**) Rasters of VMHvl^Esr1^ neuron activity and resident behavior from a representative fed (**c**) or 18-hr fasted (**d**) mouse, aligned to intruder entry. “fed” and “fasted” above the baseline raster indicate pre-trial treatment. (**e**) rSLDS model performance as cvR^2^ for *ad libitum*-fed and 18-hr fasted conditions. (**f-g**) Activity of rSLDS latent dimensions aligned with behaviors in a representative *ad libitum*-fed (**f**) and 18-hr fasted (**g**) mouse. Burgundy and gray curves indicate the X_1_ and X_2_ dimensions, respectively. (**h-i**) Behavior-triggered average (BTA) of rSLDS dimensions (X_1_-burgundy, X_2_-gray) aligned to the first sniffing bout in the fed (**h**) or 18-hr fasted (**i**) states across mice (n = 6 mice). (**j**) BTA of X_1_ dimension aligned to first attack in 18-hr fasted vs. fed states (n = 6 mice). Inset: Slope of the ramping segment from (**j_ii_**) calculated from time of first sniff. See also **Extended Data Fig. 4**d. (k) Correlation between time-to-peak of the X_1_ integration dimension and latency from first sniff to the first attack (linear regression, R² = 0.978, ****p < 0.0001). Individual mice are color-coded. (**l**) Schematic summary of results in (**j-k**) and **Extended Data Fig. 4**b-f. (**m**) BTA of X_1_ dimension aligned to the offset of the final attack bout. Dashed lines: exponential curve-fit of mean activity. (**n**) Area under the curves (AUC) in (**m**) for 100s after the last attack offset. (**o**) Time constants of rSLDS dimensions. (**p**) Dynamics of individual X_1_-weighted neurons. (**p_i_**) Stem plot of individual cell weights from 18-hr fasted mice; red circles, weights >2SD; (**p_ii_**) Activity of X_1_ -weighted neurons in fed (left) vs. 18-hr fasted (right) condition; (**p_iii_**) Representative traces of neurons from (**p_ii_**), aligned to the first sniffing bout. (**q**) ECDF of average autocorrelation half-widths (ACHWs) for X₁-weighted neurons’ activity during male-male interactions (n=6 mice). See also **Extended Data Fig. 3**g-j. (**r-s**) Representative flow fields of rSLDS models in (**f-g**) with neural trajectories color-coded by time (**r_i_, s_i_)** or behaviors (**r_ii_, s_ii_)** in both fed (**r**) and 18-hr fasted (**s**) states. *p<0.05, \*\**p*<0.01, ***p<0.001. For all figure panels, data are mean ± SEM (n=6 mice). See also **Extended Data Fig. 2**, 3 and 4.

First, we investigated whether 18hr-fasting caused changes in overall neural activity or in the behavioral tuning of individual neurons. The mean, peak z-score activity, and time to peak of z-score activity of all VMHvl^Esr1^ neurons following the first sniff were not statistically different between fasted and fed mice (Extended Data Fig. 2f-j). Choice probability analysis of single-neuron behavioral tuning^33^ confirmed that most neurons (>70%) exhibited mixed selectivity for male-directed behaviors in both the fed and 18-hr fasted states (Extended Data Fig. 2k, k, gray), consistent with prior findings in *ad libitum-* fed mice^33,48,55^. Among behavior-tuned cells, the percentage of sniff- or attack-tuned VMHvl^Esr1^ neurons was not significantly different in fed versus fasted mice (Extended Data Fig. 2k-m, q). Furthermore, the mean activity (post-sniff) of attack-tuned or sniff-tuned populations was not affected by starvation (Extended Data Fig. 2n-p, r-t). Thus, analysis of single-unit activity and behavioral tuning in VMHvl^Esr1^ neurons failed to reveal robust changes that correlated with the increased aggressiveness of 18 hr-starved mice.

We next investigated whether changes in the population coding of behavior could be observed between fed and 18hr-fasted mice. As a first step, we used generalized linear models (GLMs). Previous studies in females have indicated that GLMs using behavioral regressors do not explain a high fraction of the variance in VMHvl^Esr1^ neural activity^59^; we confirmed the same was true for activity in both fed and fasted males (Extended Data Fig. 3a, a). While inclusion of a cell-to-cell coupling term (which incorporates correlations in activity among VMHvl^Esr1^ neurons) in the GLM increased the model fit^59–61^, the mean variance in neural activity explained was still <50% in both conditions (Extended Data Fig. 3c-f), a relatively poor fit.

Because the coupling term may reflect local connectivity, and since neural dynamics can be influenced by local connectivity^62^, we next examined whether neural dynamics were altered by 18hr food deprivation. One measure of neural dynamics is their decay time following peak activity. To assess this, we calculated the autocorrelation half-width (ACHW)^33,63,64^, a proxy for single-neuron persistence, for each neuron across both conditions. This revealed a small but statistically significant increase in decay time in 18hr fasted mice in the bulk VMHvl^Esr1^ population (Extended Data Fig. 3g-j, mean ACHW: 9.7± 0.3s for fed state vs 11.58 ± 0.3s for fasted state, p****<0.0001, n=6 mice).

As VMHvl^Esr1^ neurons are heterogeneous^26^, we investigated next whether there were latent subpopulations of neurons in this population that might exhibit more robust changes in dynamics than the population as a whole, following food deprivation. To do this, we applied an unsupervised probabilistic modeling approach, recurrent Switching Linear Dynamical Systems (rSLDS)^65,66^. This approach uses dimensionality reduction to identify latent dimensions (factors) in VMHvl^Esr1^ population activity that differ in their activity and dynamics. It then fits a linear dynamical system model to activity in this latent dimension state-space, using machine learning (Extended Data Fig. 4a). Consistent with prior studies^46,59^, rSLDS models fit to data from both fed and 18 hr-fasted mice explained a much larger fraction of the variance in neural activity (cvR^2^ ≥90%; Fig. 2e) than was explained by GLMs.

To interpret these models, we examined the “learned” dynamics matrix *A* of the longest-lived state, fit by the model (Extended Data Fig. 4a). The eigenvalues of this matrix yield the time constants of each of the latent dimensions^65,66^. Consistent with our earlier studies^46–48,59^, models fit to data from fed mice yielded one dimension (*X_1_*) with a substantially greater time constant than the next-slowest dimension (*X_2_*), suggestive of a “slow mode,” or integration dimension (Fig. 2f-i, o; Extended Data Fig. 4g). This slow mode is characterized by "ramping up" behavior during sniffing episodes, reaches its peak level at the onset of attack, remains elevated between attack bouts and decays slowly thereafter^46^ (Fig. 2f-i, burgundy curves; Extended Data Fig. 4b). In contrast, the orthogonal (X_2_) dimension exhibited transient activity that peaked at the first sniff and then declined rapidly (Fig. 2f-l, gray curves; Extended Data Fig. 4c). In 2D flow-field visualizations of the state-space model, the *X_1_* dimension yielded a linear region of “slow points” that may represent a leaky integrator or approximate line attractor^46,47^ (Fig. 2r, s; Extended Data Fig. 4h, dashed lines). rSLDS models fit to data from 18 hr fasted mice yielded a qualitatively similar result, in terms of latent factors and their dynamics (Fig. 2g, i).

To determine whether line attractor parameters differed quantitatively between fed and 18 hr-fasted mice, we compared the ramping and decay phases of the projected *X_1_* integration dimension activity under the two conditions. Alignment of normalized z-scored activity to the onset of the first attack revealed significantly steeper ramping dynamics in fasted vs. fed mice (Fig. 2j_i_). Importantly, comparison of raw z-scored activity indicated that this faster ramp-up was due to a steepening of the slope rather than to a reduction in absolute peak *X_1_* activity (Fig. 2j_ii_, inset; Extended Data Fig. 4d, d_ii_). This faster ramp-up in starved mice mirrored the reduced latency from the first sniff to the first attack in this condition (Fig. 1h, I; Extended Data Fig. 4ei). Indeed, these two parameters were strongly and significantly correlated both within and across mice (Fig. 2; Extended Data Fig. 4e, e). Thus, 18-hr food deprivation caused a change in neural dynamics during the transition from sniff to aggression, not a lowering of the threshold level of *X_1_* activity associated with attack (Fig. 2l).

Our previous studies have shown that following ramp-up, activity in the line attractor remains at a stable position during attack and then slowly decays back down the attractor after the intruder is removed and fighting stops^46^. Importantly this decay constant is highly correlated with the intensity of aggressiveness across individual mice^46,48^. We therefore investigated whether the increased duration and frequency of attack in 18 hr-starved mice might be reflected in an increased stability of the line attractor. Indeed, activity in the *X_1_* dimension following the termination of attack decayed more slowly in fasted than in fed mice (Fig. 2m, n), reflecting a significant increase in both the absolute magnitude of the *X_1_* time constant, and in the *X_1_:X_2_* time constant ratio (Fig. 2o; Extended Data Fig. 4g). Consistent with this, the ACHWs of neurons strongly weighted on the *X_1_* dimension were significantly longer in fasted than in fed mice (Fig. 2p, q, mean ACHW: 11.6 ± 0.6s for fed state vs 14.9 ± 0.7s for fasted state, p***=0.0006, n=6 mice), and were greater than in the bulk VMHvl^Esr1^ population (Fig. 2q vs. Extended Data Fig. 3j). Taken together, these data indicate that the effect of 18-hr fasting to increase aggressive behavior is correlated with changes in two dynamic *X_1_* parameters: faster ramp-up and slower decay along the attractor.

In 2D flow field maps of neural state space, under both conditions the trajectory of the latent factor activity vector reached the end of the line attractor when attack commenced (Fig. 2r_ii_, s_ii_). However, the initial trajectory was different with respect to the *X_2_* dimension in fasted mice, initially following a path at an acute angle to the attractor before relaxing to the line itself, rather than moving up the attractor as in fed mice (Fig. 2r vs. 2s; Extended Data Fig. 4h). The raw activity and initial dynamics in the *X_2_* dimension appeared similar in both conditions, although there was a trend to lower *X_2_* peak activity in fasted mice (Extended Data Fig. 4c). Activity in *X_2_* may represent novelty or social fear^46^, and is known to interact with *X_1_*^47^; it is possible that the strength of this interaction is enhanced by starvation.

### Prolonged fasting reveals a biphasic effect of food deprivation on the stability of VMHvl^Esr1^ attractor dynamics

Changes in neural dynamic parameters and behavioral metrics observed between *ad libitum*-fed and 18-hr fasted mice indicate that moderate food deprivation robustly enhances both measures, consistent with the ‘hangry’ state described in humans^12,13^ and other species^7,8,9, 15^. To reconcile this with reports that starvation suppresses aggression in mice^6,14^ and with our finding that prolonged fasting (40 hr) markedly reduces aggression (Extended Data Fig. 1d), we next examined neural population dynamics under this latter condition (Fig. 3a).

**Fig. 3.**
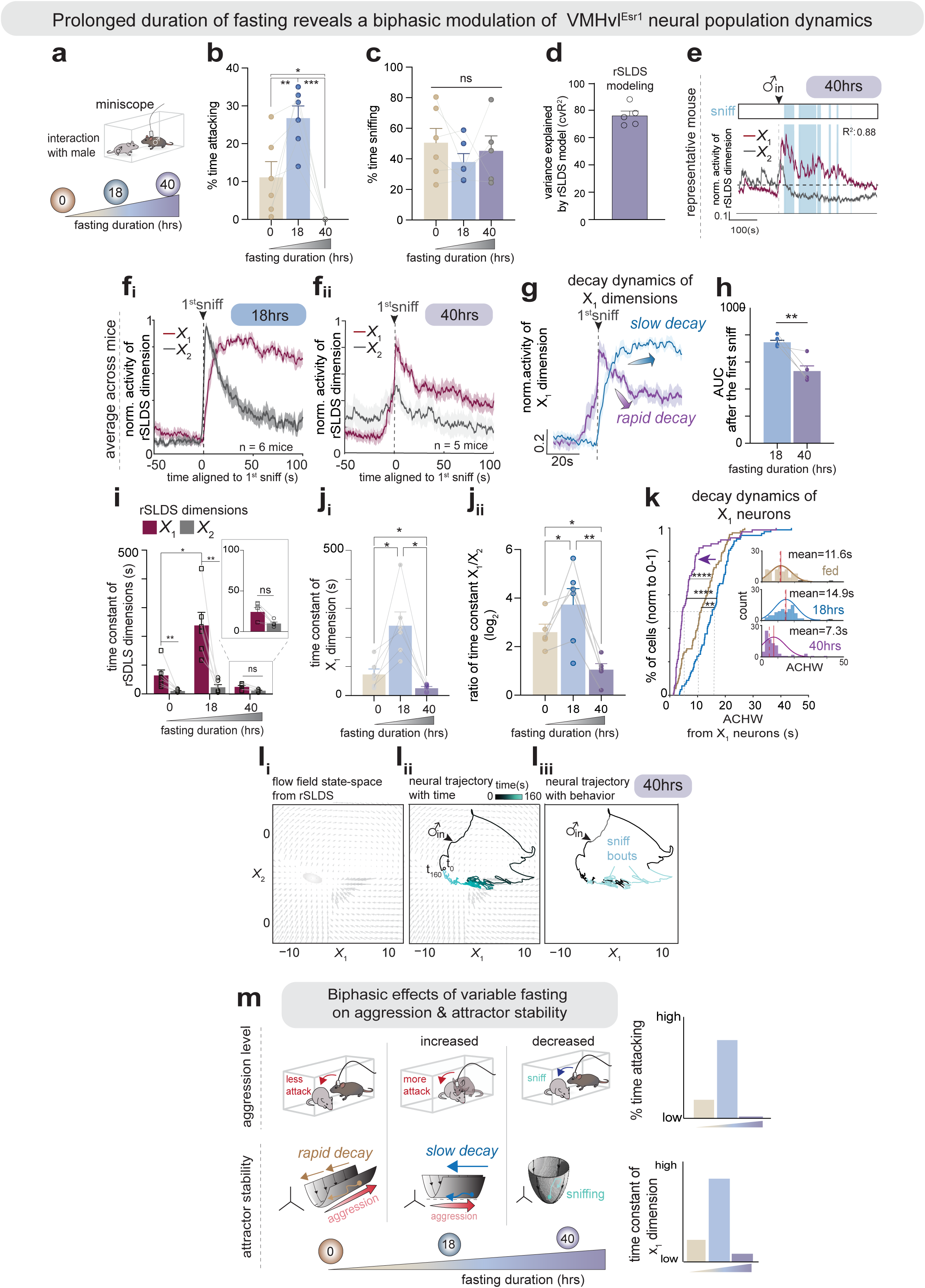
Prolonged fasting duration reveals biphasic effects on aggression and stability of VMHvl^Esr1^ attractor dynamics. (**a**) Schematic of prolonged fasting and its effects on resident-intruder assay. (**b, c**) Quantification of aggression parameters: (**b**) time spent in attack (%) and (**c**) time spent in sniff (%) across various fasting durations in mice with miniscope. See also **Extended Data Fig. 1**d, f and 2a. (**d**) rSLDS model performance as cvR^2^ for 40-hr fasted conditions. (**e**) Activity of rSLDS latent dimensions aligned with behaviors in a representative 40-hr fasted mouse. Burgundy and gray curves indicate the X_1_ and X_2_ dimensions, respectively. (**f**) BTA of rSLDS dimensions (X_1_-burgundy, X_2_-gray) aligned to the first sniffing bout in the 18-hr fasted or 40-hr fasted states across mice (n = 6 or 5 mice). **f_i_**: Replication of Fig. 2i for purposes of comparison. (**g, h**) X_1_ dynamics and activity in 40 hr-fasted mice. (**g**) BTA of X_1_ dimension aligned to the first sniffing bout in 18-hr (blue) vs. 40-hr (purple) fasted states; (**h**) AUC in (**g**) for 100s after first sniffing onset. (**i**) Time constants of rSLDS dimensions across various fasting conditions. i-inset: enlarged time constants in 40-hr fasted condition. (**j**) Comparison of time constants of X_1_ dimension (**j_i_**) or ratio of time constants from the X_1_ and X_2_ dimensions (**j_ii_**) across fasting conditions. (**k**) ECDF of ACHWs from X₁-weighted neurons’ activity during male-male interactions across fasting conditions. (**l**) Representative flow field of rSLDS models (**l_i_)** in (**e**) with neural trajectories color-coded by time (**l_ii_**) or behaviors (l**_iii_**) in a 40-hr fasted state. (**m**) Schematic summary of results in (**b-l**) illustrating the biphasic effect of varying food deprivation levels on the aggressiveness and stability of VMHvl^Esr1^ population dynamics. \**p*<0.05, \*\**p*<0.01, \*\*\**p*<0.001, \*\*\*\**p*<0.0001; ns, not significant. For all figure panels, data are mean ± SEM (n = 5 or 6 mice).

First, we confirmed that *Esr1^Cre/+^* mice implanted with a GRIN lens and miniscope responded to 40-hr starvation like wild-type unoperated mice, exhibiting a marked decrease in aggressive, but not investigatory, behaviors (Fig. 3b, c). rSLDS modeling revealed that, in contrast to the 18-hr fasted condition (Fig. 3f_i_-burgundy, g-blue), activity projected onto the *X_1_* dimension in 40-hr fasted animals exhibited rapid decay following the first sniffing bout (Fig. 3e, f_ii_-burgundy, g-purple, h). Accordingly, the time constant of the *X_1_* dimension was not significantly greater than that of *X_2_* dimension (Fig. 3i, inset), and was markedly reduced relative to both the 18-hr fasted condition (Fig. 3i, j_i_; 18-hr vs. 40-hr) and *ad libitum*-fed controls (Fig. 3j_i_; 0 vs. 40-hr). The same was true for the ratio between the time constants of *X_1_* and *X_2_* (Fig. 3j_ii_).

In agreement with these changes in latent factor dynamics, analysis of ACHWs in *X_1_*-weighted neurons identified using the emission matrix demonstrated reduced persistence and shorter decay times in 40-hr starved vs. the 18-hr fasted and fed states (Fig. 3k). Concordantly, 2D vector flow-field visualizations of rSLDS models from 40-hr fasted mice failed to exhibit a well-defined linear region of slow points (Fig. 3l, cf. Fig. 2r, s). Collectively, these data suggest that in 40-hr fasted mice, the line attractor is either absent or is considerably “leakier” than that in either *ad libitum*-fed or 18-hr fasted mice. These findings reveal a biphasic, bi-directional effect of hunger on aggression, which is reflected by a corresponding biphasic change in line attractor dynamics (Fig. 3m).

### Titrated activation of hunger-promoting neurons modulates aggression and VMHvl^Esr1^ attractor dynamics, recapitulating effects observed during naturalistic food deprivation

Our results revealed a robust interplay between starvation and aggression at the level of population dynamics in VMHvl^Esr1^ neurons. We next investigated whether the opposite-direction effects of food deprivation on aggressiveness at 18 vs. 40 hr reflected a change in the effects of a graded hunger motivational state at different intensities, or rather the gradual depletion of energy resources or weight loss after extended starvation. To distinguish between these possibilities, we tested whether acute, graded activation of hypothalamic ‘hunger’ neurons^50,54,67^ at different intensities in *ad libitum*-fed mice could recapitulate the starvation-induced biphasic effects of starvation on aggressive states and population neural dynamics.

To this end, we employed titrated chemogenetic activation of AgRP-expressing neurons in the arcuate nucleus (ARC^AgRP^ neurons); these neurons promote a hunger drive in response to interoceptive cues that signal reduced bodily energy state^50,52–54,68^. Previous studies have demonstrated that graded changes in the intensity of ARC^AgRP^ neuronal activation produced correspondingly graded increases in food intake^53,69^. Although strong activation of these neurons moderately affects aggression^6,14^, whether titrated activation can yield bi-directional effects at different intensities has not previously been investigated. To test this, we systemically administered deschloroclozapine (DCZ), a metabolically stable and fast-acting DREADD agonist, at different calibrated doses (0, [saline control], 0.1, 0.5, 1.0 mg/kg)^70^ on different days to *ad libitum-*fed AgRP^Cre/+^ mice injected with AAV-FLEX-hM3Dq-mCherry into the ARC 40 min prior to conducting RI assays (Fig. 4a; see Methods).

**Fig. 4.**
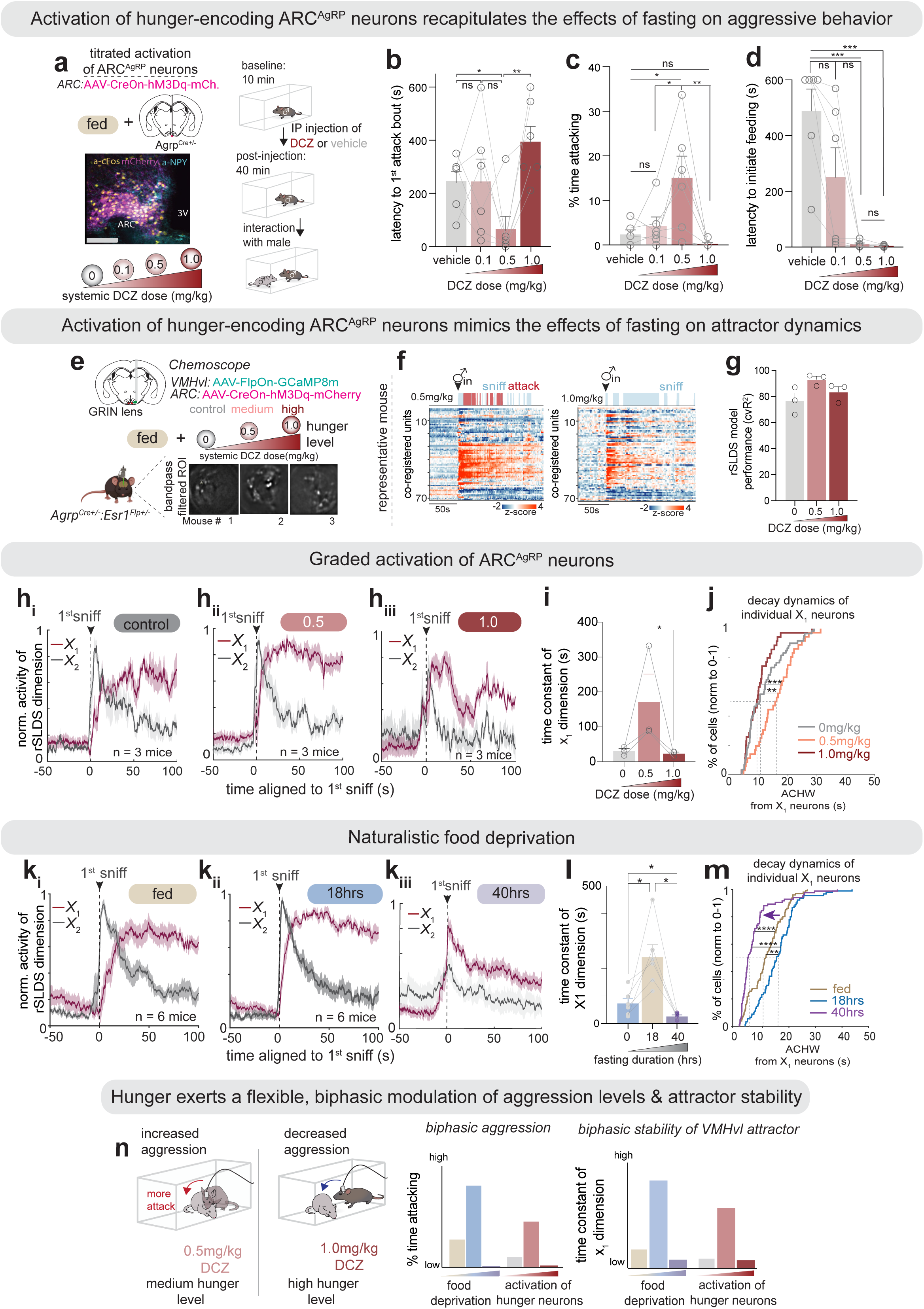
Acute, titrated activation of ARC^AgRP^ neurons phenocopies biphasic starvation-induced shifts in aggression-encoding VMH^Esr1^ attractor dynamics. (**a**) (left) Representative histological section showing hM3Dq-mCherry expression in ARC^AgRP^ neurons and cFos (yellow) induction after 1 mg/kg of DCZ treatment. (right) Schematic of activation of ARC^AgRP^ neurons by systemic DCZ doses, 40-50 min before male-male interaction in *ad libitum-*fed states. (**b-d**) Quantification of aggression parameters: (**b**) latency to the first attack, (**c**) time spent attacking (%) and (**d**) latency to the first feeding bout during titrated chemogenetic activation of ARC^AgRP^ neurons. (**e**) Titrated chemogenetic activation of ARC^AgRP^ neurons with miniscope imaging of VMHvl^Esr1^ neuronal activity. FOV from a representative mouse. (**f**) VMHvl^Esr1^ activity raster (lower, heat map) and resident behavior (upper, raster) from a representative 0.5 mg/kg (left) or 1mg/kg DCZ-treated (right) mouse, aligned to intruder entry. “0.5mg/kg” and “1mg/kg” above the baseline raster indicate pre-trial treatment (see (**a**)). (**g**) rSLDS model performance as cvR^2^ for 0, 0.5, 1.0 mg/kg DCZ conditions (n = 3 mice). (**h-j**) X_1_ dynamics with graded ARC^AgRP^ activation. (**h**) BTA of X_1_ and X_2_ dimensions, (**i**) Time constants of X_1_ dimension, and (**j**) ECDF of ACHWs of X_1_-weighted neurons under three different DCZ doses. (**k-m**) Effects of starvation. (**k**) BTA of X_1_ and X_2_ dimensions, (**l**) Time constants of X_1_ dimensions, and (**m**) EDF of ACHWs of X_1_-weighted neurons under naturalistic food deprivation conditions. Data reproduced from Fig. 2h, i**; 3f, j and k** to facilitate comparison with (**h-j**). (**n**) Schematic illustrating biphasic modulation of aggression and stability of the attractor as a function of levels of food deprivation or artificially induced hunger. \**p*<0.05, \*\**p*<0.01, \*\*\**p*<0.001, \*\*\*\**p*<0.0001; ns, not significant. For all figure panels, data are mean ± SEM. See also **Extended Data Figures 5 and 6**.

At the lowest dose of DCZ (0.1 mg/kg), there was no significant effect on attack-related parameters compared to the control condition; however, there was a marginal reduction in the latency to initiate feeding, indicating the efficacy of chemogenetic activation of ARC^AgRP^ neurons even at this relatively low dose (Fig. 4b-d, 0.1mg/kg). In contrast, at an intermediate dose (0.5 mg/kg), mice displayed a significantly shorter latency to the first attack as well as a marked increase in the time spent attacking (Fig. 4b, c, 0.5mg/kg), resembling “hangry” mice. The latency to initiate feeding was further reduced, resulting in virtually immediate engagement in feeding behavior upon food presentation in the absence of an intruder male (Fig. 4d, 0.5 mg/kg). In contrast, at the highest dose of DCZ tested (1 mg/kg), aggressive behaviors were markedly suppressed, and animals exhibited an immediate initiation of feeding upon food presentation (Fig. 4b-d, 1 mg/kg). Thus, titrated chemogenetic activation of ARC^AgRP^ neurons recapitulated the biphasic, bi-directional effect of progressively increased durations of food deprivation on aggression (Fig. 4b, c; see also Fig. 3b and Extended Data Fig. 1d).

Next, to assess whether and how the graded activation of ARC^AgRP^ neurons modulates attractor dynamics in VMHvl^Esr1^ neurons, we combined miniscope imaging of VMHvl^Esr1^ neurons with titrated chemogenetic activation of ARC^AgRP^ neurons (Fig. 4e, f). Specifically, we expressed a Flp-dependent GCaMP8m calcium indicator in VMHvl^Esr1^ neurons and a Cre-dependent FLEX-hM3Dq-mCherry virus in ARC^AgRP^ neurons of AgRP^Cre/+^; Esr1^Flp/+^ double-transgenic mice (Fig. 4e). VMHvl^Esr1^ neural activity was longitudinally recorded across multiple days under three experimental conditions: mice were subjected to systemic administration of saline (control), 0.5mg/kg (medium), and 1mg/kg (high) doses of DCZ in a physiologically *ad libitum*-fed state, using a counter-balanced design across conditions (Fig. 4f), and calcium signals were acquired during an RI assay 40 minutes following drug administration. Social behaviors were measured in synchronized video recordings (Extended Data Fig. 5a, blue and red rasters).

rSLDS models fit to VMHvl^Esr1^ neural population activity recorded at different DCZ doses yielded consistently high cvR^2^ values across all tested conditions (Fig. 4g). Notably, quantitative changes in dynamic parameters of the two latent dimensions revealed by rSLDS modeling during titrated activation of ARC^AgRP^ neurons closely mirrored the biphasic dynamical changes observed under naturalistic food-deprivation conditions (Fig. 4h-i vs. 4k-l). Specifically, under control (saline-injected) conditions, activity along the *X_1_* dimension progressively ramped up before the initiation of the first attack (Fig. 4h_i_; Extended Data Fig. 5a-left, 5c-gray), similar to *ad libitum*-fed un-injected control mice (Fig. 4k_i_). Activation of ARC^AgRP^ neurons at the medium DCZ dose, when mice showed the shortest latency to attack behavior (Fig. 4b; 0.5 mg/kg), yielded more rapid ramping followed by sustained activity in the X_1_ dimension (Fig. 4h_ii_-burgundy; Extended Data Fig. 5a-middle, 5b, c-d (pink), 5d-e), resembling the *X_1_* activity pattern in the 18-hr fasted condition (Fig. 4k_ii_). Aligning neural trajectories along the *X_1_* dimension to the onset of the initial sniff revealed a significantly accelerated ramping rate in the medium-dose condition compared to controls (Extended Data Fig. 5a; left vs. middle and 5c-e). Remarkably, the latency to attack was almost perfectly correlated across mice with the time-to-peak of projected *X_1_* activity in the medium-dose condition (Extended Data Fig. 4f; R^2^ = 0.98, p*=0.045). This correlation was even stronger than that observed under 18 hour of food deprivation (R^2^ = 0.85; Fig. 2k; Extended Data Fig. 4f-left), perhaps reflecting the more precise control over hunger levels achieved by chemogenetic activation of ARC^AgRP^ neurons.

In contrast, during chemogenetic activation of ARC^AgRP^ neurons at the highest dose tested (1 mg/kg), rSLDS analysis revealed a pronounced reduction in the persistence of activity along the *X_1_* dimension, reflected in a rapid decay following initial sniff (Fig. 4h_iii_-burgundy; Extended Data Fig. 5a-right, 5g-dark red). This rapid decay resembled the faster dynamics observed in the prolonged starvation condition (Fig. 4k_iii_), and was reflected by a reduced magnitude of the *X_1_* time constant remarkably similar to that seen under 40 hr starvation (Fig. 4i, l; Extended Data Fig. 5l). Notably, both the persistence of X_1_ activity, quantified as the area under the curve (AUC) following the initial sniff, and the ACHWs of *X_1_*-weighted neurons, were significantly diminished in the high-dose DCZ condition, compared to the medium-dose (0.5 mg/kg) condition (Fig.4j; Extended Data Fig. 5g, g, k). These dose-dependent biphasic effects were further underscored by changes in the ratio of the *X_1_* to *X_2_* time constants (Extended Data Fig. 5i, i, m), highlighting a graded, biphasic modulation of attractor stability by scaled hunger levels.

Consistent with the observed DCZ dose-dependent changes in these fit rSLDS model parameters, neural trajectories visualized in a two-dimensional flow-field state space under the medium-dose condition closely resembled those observed after 18 hours of fasting, with prominent line attractors in both conditions (Extended Data Fig. 6a vs. 6b). In contrast, the trajectories under the high-dose condition exhibited disrupted attractor dynamics reminiscent of those following 40 hours of fasting, characterized by a loss of clearly defined linear regions of slow points (Extended Data Fig. 6c vs. 6d). Changes in the ACHWs of individual *X_1_* dimension-weighted neurons under titrated chemogenetic AgRP neuron activation similarly paralleled the changes observed under naturalistic food deprivation states (Fig. 4j vs. 4m).

Thus, graded activation of ARC^AgRP^ neurons in *ad libitum*-fed mice reproduced the bi-phasic changes in both aggressive behavior and the dynamics of VMHvl^Esr1^ neuronal activity observed with different durations of food deprivation (Fig. 4n; Extended Data Fig. 6e). These data argue against the idea that the reduced aggression observed after 40 hrs of starvation is due simply to energy depletion or weight loss and imply that increasing intensities of a unitary starvation state are converted into opposite-direction modulation of aggressiveness and line attractor properties at distinct thresholds. They further suggest that the VMHvl line attractor constitutes a key integrative node where metabolic and motivational signals converge to regulate aggressiveness, likely through state-dependent modulation of underlying population dynamics. While we cannot formally exclude that hunger-induced changes in line attractor dynamics might be a consequence rather than a cause of changes in aggressive behavior, this seems unlikely given that the line attractor is present in mice passively observing, as well as actively engaging in, attack behavior^47^, and that it persists in ‘hangry’ mice in which attack is interrupted by feeding (see below).

### Increased line attractor stability in ‘hangry’ mice maintains aggressiveness during feeding

The above results establish that a graded hunger state can either enhance or suppress aggressiveness and line attractor dynamics, depending on its intensity. Given that aggression is energetically costly^71^ and mutually exclusive with feeding, these findings prompted us to ask two related questions: ‘Why does an intermediate level of hunger elevate aggression?’, and ‘Why is this increase in aggressiveness encoded in the enhanced stability of a line attractor, rather than simply by an increased activity in aggression-promoting neurons?’ To tackle these questions, we further examined attractor dynamics under conditions in which 18-hr fasted animals were allowed to choose between feeding and attacking an intruder. To this end, following an approximately 2-min RI test to assess the resident’s baseline level of aggressiveness, we introduced a chow pellet without removing the intruder (Fig. 5a). After imaging neural activity and observing behavior for two minutes in the presence of both the food and the intruder, we removed the intruder but left the food pellet in place (to determine whether the resident was satiated or still hungry). This design allowed us to examine the animal’s behavioral choices under conditions of competing hunger and aggressive drive states, with targets of both consummatory behaviors concurrently provided.

**Fig. 5.**
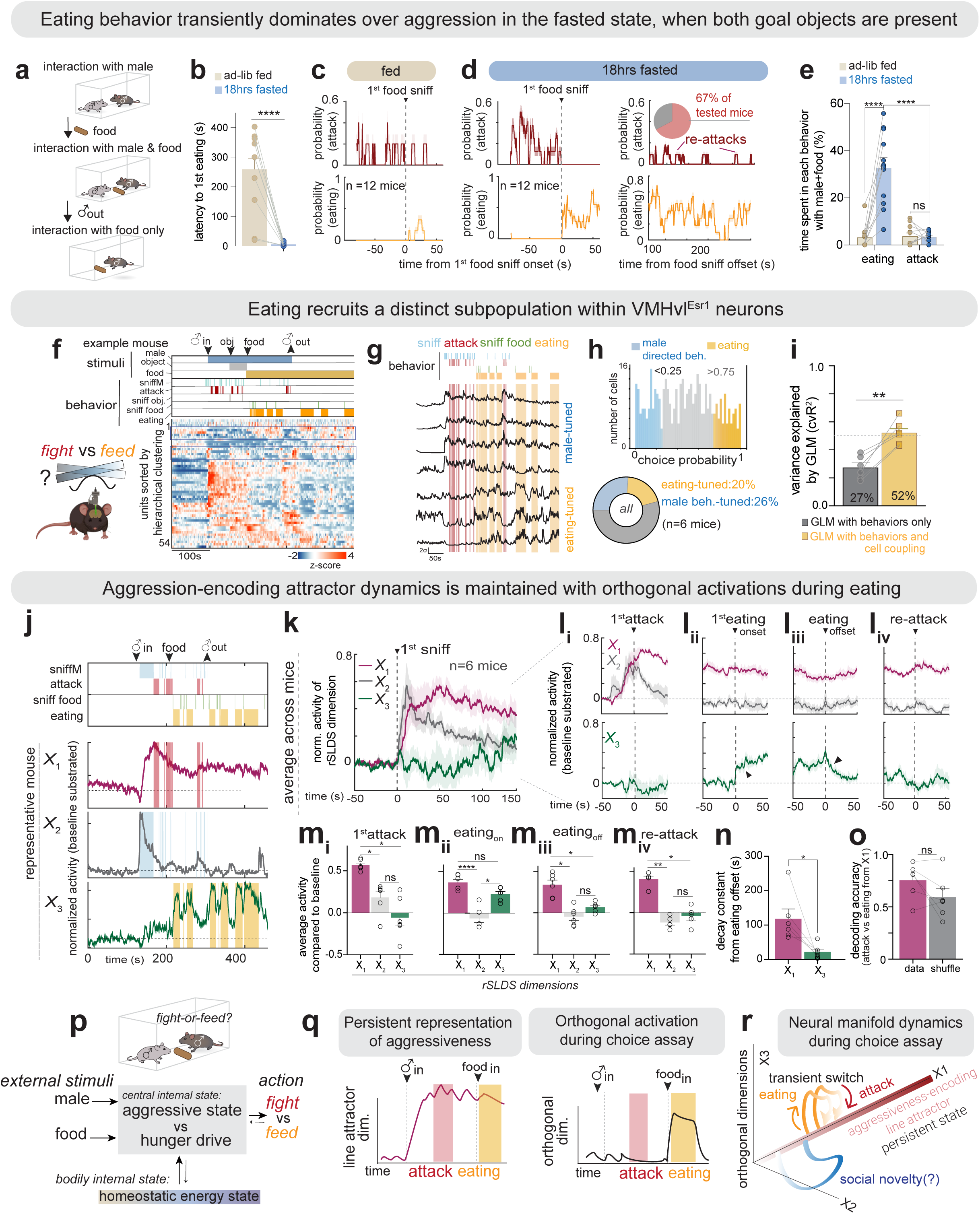
Persistent aggressiveness during intermittent fighting and feeding in hangry mice. (**a**) Schematic illustrating experimental design. (**b**) Latency to the first eating bout upon the presentation of a chow pellet in fed vs. 18-hr fasted states. (**c-d**) Probability of attack (upper) and eating behavior (lower) aligned with first food sniff onset in fed (**c**) and 18-hr fasted states (**d**-left), and aligned with the offset of first food sniff (**d**-right) (n=12 mice). (**e**) Time spent in eating and attack behaviors during the simultaneous presence of a male and food in fed vs. 18-hr fasted states. (**f**) Rasters of hierarchically clustered VMHvl^Esr1^ neuron activity (lower, heat map) with a behavior raster (upper) from a representative 8-hr fasted mouse. (**g**) Representative traces of activity from male-behavior or eating-tuned neurons in (**f**). Colored vertical bars indicate behavior raster. (**h**) Choice probability for male-directed behavior- vs. eating behavior-tuned cells (upper) and pie chart showing proportion of tuned cells (lower) (n=6 mice). (**i**) Quantification of cvR^2^ for GLM with male or food-related behaviors only vs. behaviors with a cell-coupling term from neural activity (n = 6 mice). See also **Extended Data Fig. 7**c. (**j**) (lower) First three dimensions of an rSLDS model (X₁-X₃) from a representative mouse, aligned to male entry; (upper) behavior rasters. (**k**) BTA of rSLDS dimensions X₁ (burgundy), X₂ (gray), and X₃ (green) dimensions aligned to the first sniff bout (n = 6 mice). (**l**) BTAs of baseline-subtracted, normalized activity of X₁ and X₂ (upper row) and X₃ (lower row) aligned to (**l_i_**) first attack, (**l_ii_**) first eating bout, (**l_iii_**) eating offset, and (**l_iv_**) re-attack. (**m**) Quantification of BTAs in X₁–X₃ dimensions during the four epochs in (**l**). One-way ANOVA with paired post-hoc tests with multiple comparison correction. (**n**) Empirically measured decay constants of X₁ vs. X₃ activity following eating offset (**l_iii_**). (**o**) Accuracy of decoding eating vs. attack bouts using X_1_ dimension activity (data) is not significantly better than shuffled control, reflecting similar level of X_1_ activity during both behaviors (**l_i,ii_**) (n = 6 mice). (**p-r)** Schematics summarizing design (**p**) and results (**q, r**) from behavioral choice assay. (**q**) Summary of activity in X_1_ (left) vs. X_3_ (right) dimensions during attack vs. eating. (**r**) Neural state-space trajectories of activity in rSLDS latent dimensions during periods of attacking vs. eating. *\*p<*0.05*, **p<*0.01*, ****p<*0.0001; ns, not significant. For all figure panels, data are mean ± SEM. See also **Extended Data Figures 7 and 8**.

Following the introduction of the chow pellet, ‘hangry’ mice immediately stopped fighting, investigated the pellet, and ate (Fig. 5b, d-left). In contrast, relatively little feeding was observed in *ad libitum* fed animals, as the chow pellet only yielded brief interruptions of attack (Fig. 5b, c). In the 18-hr fasted mice, following a period of feeding (∼100 s), 67% of the fasted mice resumed attacking the intruder (Fig. 5d-right; Extended Data Fig. 7a, b-pie chart). However, their latency to resume attack was marginally longer than in the *ad libitum*-fed control condition (mean latency to resume attack after the offset of the first food sniff (s): 137 ± 26 s (fasted) vs. 94 ± 24 s (fed), p=0.06) (Fig. 5d; Extended Data Fig. 7b). Following the initial re-attack, 18-hr fasted mice resumed feeding and continued with interspersed feeding and attack bouts (Fig. 5d-right; Extended Data Fig. 7a). Overall, the fasted mice spent far more time feeding than attacking (percent time spent eating vs. attacking in fasted state: 32.7 ± 4.42 % (eating) vs. 3.37 ± 0.67 % (attacking), ****p<0.001; in fed state: 3.2 ± 3.7 % (eating) vs. 3.7 ± 1.3 % (attacking), p=0.882, Fig. 5e; Extended Data Fig. 7a-right). Nevertheless, under these conflicting conditions mice did not achieve satiety, as indicated by extensive feeding following removal of the intruder (Fig. 5f, yellow raster). These findings reveal a dynamic and hierarchical interplay between two competing motivational states and suggest that mice are willing to delay or sacrifice full repletion of their energy reserves in order to defend their food resource from a competitor male.

We next sought to determine how neural activity and dynamics in VMHvl^Esr1^ neurons might be modulated during the observed dynamic switching between fighting and feeding. Unexpectedly, hierarchical clustering as well as choice probability analysis of single VMHvl^Esr1^ neuron activity revealed a robust population of eating-tuned neurons in 18hr fasted mice (Fig. 5f-h). Consistent with this observation, a larger fraction of variance in neuronal activity could be explained by a behavior-weighted GLM in fasted mice during interactions with food and a male, compared to GLMs fit to the same mice in the absence of food but presence of male (mean cvR^2^ in fasted mice with food: 27% variance explained, Extended Data Fig. 7ci; mean cvR^2^ in fasted mice without food: 17% of variance explained, Extended Data Fig. 3b, lower), likely reflecting the behavior-locked activity of the eating-tuned population (see below; Fig. 5h). Nevertheless, the addition of a cell-coupling term still increased the variance explained by GLMs in the presence of food (mean cvR^2^: 0.27 with only behavioral regressors, cvR^2^: 0.52 with behaviors and cell-coupling, Fig. 5i; Extended Data Fig. 7cii). This suggests that local functional connectivity between neurons - and therefore potentially neural dynamics - influences population activity in this paradigm as well.

To directly examine neural dynamics in this competition paradigm, we fit rSLDS models to VMHvl^Esr1^ activity. In this condition, models with three dimensions instead of two explained a greater fraction of neural variance in data (66.3 ± 4% variance explained for the 3D model vs. 44.1 ± 2% variance explained for the 2D model, Extended Data Fig. 7d). To understand the behaviors represented by these three latent factors, we analyzed their projected time-varying activity during sniffing, attack, and eating (Fig. 5j; Extended Data Fig. 7e, f). The *X_1_* dimension exhibited ramping activity that peaked near the attack onset and persisted thereafter (Fig. 5j, k, l_i_, m_i_-burgundy; Extended Data 7e, f, *X_1_*), as previously described^46^. Interestingly, despite the switch of overt behavior from attacking to eating, the level of activity in the *X_1_* dimension remained stable and elevated during food sniffing and consumption (Fig. 5j, k, l_ii__-iii_, m_ii-iii_, burgundy; Extended Data Fig. 7e, f, X_1_). Notably, when animals resumed attacking, activity in the X_1_ dimension was similar to that observed during the initial attack (Fig. 5l_iv_, m_iv_; Extended Data Fig. 7e, f, X_1_ and 7i). By contrast, activity in the *X_2_* dimension declined following the initial male interaction and remained relatively low throughout aggression, feeding and re-attack (Fig. 5j, k, l_i-iv_, m_i-iv_-gray; Extended Data Fig. 7e, f, X_2_).

In contrast to these first two dimensions, the *X_3_* dimension, enriched in food-modulated neurons (Fig. 5j, X_3_), displayed little activity during male interactions before the delivery of the food pellet, but exhibited time-locked, transient increases in activity during feeding bouts (Fig. 5l_ii_-arrowhead, 5m_ii_-green; Extended Data Fig. 7e, f, X_3_). Following the offset of eating, the *X_3_* dimension displayed rapid decay, unlike the slow decay observed in the *X_1_* dimension (Fig. 5l_iii_-arrowhead, n), reflecting distinct dynamics across these two dimensions of activity.

These results indicate that activity in the *X_1_* line attractor-aligned dimension remained stably elevated as the animal switched back and forth between attacking and eating (Extended Data Fig. 7g). Indeed, a decoder trained on the activity of the *X_1_* dimension was unable to distinguish video frames containing attack vs. eating (Fig. 5o), whereas a decoder trained on the activity of the *X_3_* dimension robustly distinguished attack vs eating frames (Extended Data Fig. 7h). In line with this population-level dissociation, neurons with strong weights onto the *X_3_* dimension exhibited significant tuning for eating, whereas neurons contributing to the *X_1_* and *X_2_* dimensions were preferentially tuned to male-directed behaviors (Extended Data Fig. 7i, j).

To visualize the dynamics of these three dimensions in neural state space, we first plotted the fit rSLDS state-space model as a vector flow field spanned by the *X_1_-X_2_* dimensions. In this space, we observed that the population vector remained in the line attractor during bouts of eating as well as attack (Extended Data Fig. 8a-d). In contrast, the fit vector flow field spanned by the *X_1_-X_3_* dimensions revealed that the population vector displayed movements orthogonal to the line attractor and parallel to the *X_3_* dimension during eating but returned to the line attractor immediately when the animals switched back to attack (Extended Data Fig. 8e-h).

Together, these findings revealed that neural population dynamics in the VMHvl^Esr1^ neurons of mice in a behavioral choice paradigm exhibited a persistently elevated component of activity in the *X_1_* (line attractor) dimension, despite repeated switches between attack and eating. Transitions to eating engaged an orthogonal dimension (*X_3_*), which (in contrast to *X_1_*) displayed rapid decay following the offset of feeding. As attack resumed following such feeding bouts, activity in the line attractor dimension dominated over that in the orthogonal dimensions (*X_2_* and *X_3_*) (Fig. 5p-r).

## DISCUSSION

While VMHvl is well-known to regulate both social behaviors^20,22,72^ and feeding^73,74^, relatively few studies have investigated how these two functions influence each other^21^ and none has done so at the level of neural population activity. Here we have investigated how aggressiveness and neural dynamics are influenced by hunger in VMHvl. We find that starvation exerts bi-phasic, opposite-direction effects on aggression depending on the duration of food deprivation^6,14,15^, and that these bi-phasic effects are reflected in bi-phasic changes in line attractor dynamics^46^ rather than in overall levels of activity among VMHvl^Esr1^ neurons. For example, our analysis suggests that ‘hangry’ mice do not have a “shorter fuse” than fed mice, as intuition might suggest; rather, their “fuse” is the same length, but burns faster (increased ramping rate from baseline to peak in the *X_1_* dimension). Together, our observations reveal how population neural dynamics that encode aggressiveness, an affective or emotional internal state^75^, can be modified by a competing homeostatic physiological drive state^76,77^.

Importantly, the bi-phasic effects of food deprivation on aggressiveness and line attractor dynamics could be replicated by acute, titrated chemogenetic activation of ARC^AgRP^ neurons^52,53^ in sated mice. This finding argues that these bi-directional effects reflect different intensities of a graded hunger state^5,78^, and that the loss of aggressiveness after 40 hrs of starvation^5,6,14^ is not due simply to gradual depletion of energy reserves or to weight-loss. These observations make sense from an ecological perspective^79^. In this view, the level of risk the animal is willing to tolerate scales with the intensity of its starvation state: if moderately hungry, the animal will fight to protect its food supply, risking further energy depletion (and injury or death if defeated). With extended starvation, however, the animal prioritizes food consumption, during which time it is vulnerable to attack. The circuit-level mechanisms whereby continuous increases in the level of hunger cause opposite-direction effects on aggressiveness at different thresholds are likely complex^21,80^, and remain to be investigated.

Why should the competition between hunger and aggressiveness be reflected by changes in line attractor dynamics? An attractor provides a region of stability in neural state space, to which neural activity rapidly returns if subjected to an off-manifold perturbation^40^. When “hangry” animals engaged in fighting were presented with food pellets, they initially switched from fighting to feeding. Although neural population activity moved out of the attractor during food consumption, activity in the attractor dimension remained elevated. Following such feeding bouts, activity rapidly returned to the line attractor dimension as attack resumed (Fig. 5r). These observations suggest a potential adaptive role for the VMHvl line attractor, namely, to keep hungry animals primed for aggression while feeding in the presence of an intruder. At the manifold level^37,38^, we speculate that the “attractive” nature of the integration dimension may provide a counterpoint to the strong motivation to feed, thereby mediating a delicate balance between these competing drives. How this balance is computed and read out to control behavior will be an important question for future investigation.

## Acknowledgements

We thank J.S. Chang, X. Da, X. Wang, D. Wagenaar, M. Yee, and L. Salay for technical assistance, K. Lee for mouse care, G. Mountoufaris and K. Cheung for training in miniscope imaging, S.W. Linderman and members of the Anderson laboratory for helpful discussions, L. Chavarria for administrative assistance, and C. Chiu for lab management. This work was supported by NIH grants R01NS123916 and R01MH123612, and the Simons Foundation to D.J.A., a HHMI-HHWF postdoctoral fellowship and postdoc fellowship abroad from the National Research Foundation of Korea (NRF) to J.K., and a National Science Scholarship from the Agency of Science, Technology and Research, Singapore and the Nanyang Assistant Professorship Start-up Grant from Nanyang Technological University, Singapore to A.N. D.J.A. is an Investigator of the Howard Hughes Medical Institute.

## Author contribution

J.K. and D.J.A. conceived the project. J.K. performed all experimental work with help from S.H., A.V., and M.L. J.K. and A.N. performed computational analyses. N.C. and A.N. developed data analysis pipelines with help from J.X. J.K., A.N., and D.J.A. analyzed and interpreted the data and co-wrote the manuscript. D.J.A. supervised the project.

## Competing interests

The authors declare no competing interests.

## METHODS

### Animals

All experimental procedures involving the use of live mice or their tissues were carried out following NIH guidelines and approved by the Institute Animal Care and Use Committee (IACUC) and the Institute Biosafety Committee (IBC) at the California Institute of Technology (Caltech). All mice in this study, including wild-type and transgenic mice, were bred at Caltech or purchased from Charles River Laboratory. Group housed C57BL/6N female or singly housed male mice (2-5 months) were used as experimental mice. *Esr1^cre^* mice, *Esr1^flp^* mice were backcrossed into the C57BL/6N background (>N10) and bred at Caltech. Heterozygous *Esr1^cre^* or double heterozygote *Esr1^flp/+^AgRP^cre/+^* mice were used for cell-specific targeting experiments and were genotyped by PCR analysis using genomic DNA from tail tissue. All mice were housed in ventilated micro-isolator cages in a temperature-controlled environment (median temperature 23°C, humidity 60%), under a reversed 11-h dark–13-h light cycle, with *ad libitum* access to food and water. Mouse cages were changed weekly.

### Housing conditions for social and sexual experience

All male C57BL/6N mice used in this study were socially and sexually experienced. Mice aged 8–12 weeks were initially co-housed with a female BALB/c mouse for 1 week and were screened for male-typical social behaviors. Mice that showed both mounting towards females and attacking towards males during a 30-minute resident intruder assay were selected for surgery and subsequent behavior experiments. From this point forward, these male mice were always co-housed with non-pregnant female BALB/c mice.

### Stereotaxic Surgeries

Virus injection and implantation were performed as described previously^33,37^. Briefly, animals were anesthetized with isoflurane (5% for induction and 1.5% for maintenance) and placed on a stereotaxic frame (David Kopf Instruments). The virus was injected into the target area using a pulled-glass capillary (World Precision Instruments) and a pressure injector (Micro4 controller, World Precision Instruments), at a 20 nl min^−1^ flow rate. The glass capillary was left in place for ten minutes following injection before withdrawal. Integrated GRIN Lenses (Inscopix) were slowly lowered into the brain and fixed to the skull with dental cement (Metabond, Parkell). After virus injection and lens implantation, males were co-housed with an ovariectomized or non-pregnant female mouse. Four to six weeks after lens implantation, mice were head-fixed on a running wheel, and a miniaturized micro-endoscope (nVista, Inscopix) was lowered over the implanted lens until GCaMP-expressing fluorescent neurons were in focus.

### Virus injection and GRIN lens implantation

The following AAVs were used in this study, with injection titers as indicated. Viruses with a high original titer were diluted with clean PBS on the day of use. AAV1-EF1a-FLEX-GCaMP8m (∼2.3 × 10^12^vg/ml, Addgene), AAVdj-hSyn-fDIO-GCaMP8m (∼2.0×10^12^ vg/ml, Janelia Vector Core), and AAV8-DIO-hM3Dq (4×10^12^ vg/ml, Addgene).

#### Stereotaxic coordinates for the viral injection (unless specified)

VMHvl, AP: −1.6, ML: ±0.78, DV: −5.7 (from the skull surface)
ARC, AP: −1.7, ML: ±0.25, DV: −5.85 mm (from the bregma)

Stereotaxic injection coordinates were based on the Paxinos and Franklin atlas. GRIN lens implantation: VMHvl, AP: −1.6, ML: −0.76, DV: −5.55 (⌀0.6 × 7.3 mm GRIN lens).

### Behavioral assays

Prior to resident-intruder (RI) assays, male mice were co-housed with females to have sexual experience for at least one week. Before the assay, cohoused females were removed from the home cage for at least two days. Male mice were acclimated to the behavioral room with a dimmed red light for two consecutive days, followed by a third day of habituation for five minutes of interaction with a male Balb/c mouse to reduce any novelty. The day before the RI assay, mice were separated into ‘fed’ or ‘fasted’ cohorts, counter-balanced for the order of conditions. At the onset of fasting (food removal), every mouse in the relevant cohort was singly transferred into a new home cage with fresh bedding. On the day of the RI assay, mice were habituated to the behavior chamber for 10 minutes, followed by a ‘baseline’ for 5 minutes. A novel Balb/c male (or for mating assays, a sexually experienced/hormonally primed female mouse) was introduced, and the RI session continued for 5 or 10 minutes. For the choice assay, following an initial 2 min interaction with a male intruder either a 2.5-4g normal chow pellet (or, as a control, the cap of a 15ml Falcon tube) was dropped into the home cage, leaving the intruder in place. After two minutes of observation, the intruder was removed and the test mouse left alone with the food pellet for an additional 5-10 minutes to assess its feeding behavior.

### Behavior recording

All behavioral experiments were performed in conventional mouse housing cages (the resident mouse’s home cage) under red lighting, using the previously described behavior recording setup ^81^. The behavior video’s top and front views were acquired at 30 Hz using StreamPix9 (Norpix) video recording software.

### Behavior Annotations

Behavior videos (top and side views) were manually annotated using a custom MATLAB-based behavioral annotation interface called BENTO^57^.

We manually annotated the following resident male behaviors:

During male-male interactions: sniffing the male (body, genital, and face), dominance mounting, and attack. For analysis of the imaging experiment, we also detailed the intruders’ behaviors, including exploration of the cage, immobility, and defensive postures (exposing their abdominal parts while standing). During male-female interactions, sniffing the female (body, genitals, and face), mounting, and intromission were scored. During male-food interaction, sniffing food and eating were scored.

### Chemogenetic activation

Behavioural tests were performed at three day intervals, at minimum. The order and number of mice receiving saline or DCZ (Sigma) were counterbalanced across different conditions. DCZ was dissolved in DMSO and further diluted in saline; DCZ (0.1, 0.5, 1.0 mg/kg of body weight) or saline (control) was intraperitoneally (i.p.) injected 40-50 min before behavioral tests.

### Chemoscope imaging

For experiments combining miniscope imaging with chemogenetic manipulations (“Chemoscope”^82^), to minimize population activity changes associated with stress from i.p. injections, a series of habituation sessions was performed every two consecutive days before any test conditions used for analysis. Mice were injected with 10ul/g (body weight) of saline (control) and returned to their home cage for 40-50 min before imaging^82^. On the test day, mice were injected with control or 10ul/g of different doses of DCZ (0.5 vs 1.0mg/kg). The order and number of mice in each condition were randomized, and other conditions were performed at 2-3 days intervals. On the day of imaging, mice were injected 40-50 mins before the RI assay.

### Miniscope data acquisition and extraction

On the day of imaging, mice were habituated for at least 10 minutes after installing the miniscope (Bruker, Inc.) in their home cage before the start of the behavior tests. Imaging data were acquired at 30 Hz with spatial down sampling (2); light-emitting diode power (0.2) and gain (3–6) were adjusted depending on the brightness of GCaMP expression as determined by the image histogram according to the user manual. A transistor– transistor logic (TTL) pulse from the Sync port of the data acquisition box (DAQ, Inscopix) was used for synchronous triggering of StreamPix9 (Norpix) for video recording. Imaging sessions typically lasted 30 mins per day. Preprocessing and motion correction were performed using Inscopix Data Processing Software. Briefly, the raw imaging data were spatially sampled twice, motion corrected, and temporally downsampled to 10 Hz, and then exported as a .tiff image stack. A spatial band-pass filter was then applied to remove the out-of-focus background. After preprocessing, calcium traces were extracted and deconvolved using the CNMF-E large data pipeline^83^ with the following parameters: patch_dims = [32, 32], gSig = 3, gSiz = 13, ring_radius = 15, min_corr = 0.7-0.8, min_pnr = 8. Every extracted unit’s spatial and temporal components were carefully inspected manually (signal-to-noise ratio, peak-to-noise ratio, size, motion artifacts, decay kinetics, and so on), and outliers (apparent deviations from the normal distribution) were discarded. The extracted traces were then z-scored before further analysis.

### Longitudinal imaging data extraction and preprocessing

We imaged the male mice implanted with a GRIN lens for multiple days under varying durations of food deprivation in a non-consecutive manner with two to three days’ intervals. On the day of imaging, we compared the field of view with a reference mark to ensure the accuracy of registration of the spatial FOV across different days. Miniscope data was selected and concatenated into one .isxd file. Data was preprocessed, and the traces were extracted as described in the last section. The extracted traces were z-scored and split into multiple traces for individual days^59^. If two imaging sessions were compared directly, the cell registration algorithm^83^ in the CIAtah was used to extract traces in the registered cells.

#### Immunohistochemistry

For Fig.4a, animals were anesthetized with isoflurane 90 mins after 1 mg/kg DCZ treatment and cordially perfused with 30 ml of cold 4% PFA following 30 ml of 0.9% saline. Brains were post-fixated for 4 hrs in 4% PFA at 4°C and then transferred to 30% sucrose in PBS. Brains were coronally sectioned at 60 μm using a Leica cryostat. The sectioned brains were washed with PBS for 10 mins and blocked with 2 % normal donkey serum in 0.2% PBST for 1 hr at R.T. Primary antibodies, goat anti-cFos (1:200, Santa Cruz, sc 52-g), mouse anti-NPY (1:1000, Santa Cruz, sc-133080), in the blocking solution (2% NDS in 0.2% PBST-Triton X-100) were incubated at 4°C for 72 hours. The sections were washed with PBS (3 x 15 mins), followed by incubation of the second antibody, donkey anti-mouse Alexa 647 (1:500, Life Technologies, A21207) or donkey anti-goat Alexa 488 (1:500, Life Technologies, A11055), for 1 hr at RT. Then they were washed with PBS (2 x 15 mins), stained with DAPI (1:10000) for 6 mins, and mounted with DAKO mounting medium. Confocal images were captured with an Olympus microscope.

### Data Analysis

#### Calculation of pixel changes and pose estimation using MARS

Pose estimation was performed using MARS^57^ to identify specific body parts ("keypoints") of the resident mouse and the intruder mouse during behavioral assays. After identifying keypoints, pixel changes were computed to quantify pose dynamics during behavior bouts, such as sniffing or attack. Specifically, pixel change was calculated within a defined region—a square measuring 20×20 pixels—surrounding each identified keypoint on the resident mouse. This calculation was performed on a frame-by-frame basis throughout the duration of each bout of sniffing or attack behavior. This metric of pixel change thus provides a quantitative measure of pose dynamics and movement intensity localized to each relevant body part, allowing comparisons of these parameters between fed and fasted conditions.

### Choice Probability (CP)

Choice probability (CP) analysis was performed as previously described^33,55^ to quantify a neuron’s selectivity or “tuning,” defined here as the ability to discriminate between two behavioral conditions based on single-cell activity^84^. For each neuron, the CP was calculated by constructing histograms of its ΔF(t)/F0 activity values during the two conditions of interest. These histograms were compared to generate a receiver operating characteristic (ROC) curve, from which CP was determined by calculating the integral of the area under the curve (AUC). CP values range from 0 to 1, with 0.5 representing chance-level discrimination. To assess statistical significance, we performed a shuffle-based permutation test by randomizing the timing of behavioral events across the two conditions and recalculating CP 100 times. Observed CP values were considered significant if they exceeded two standard deviations (2σ) above the mean shuffled CP distribution and additionally surpassed an absolute CP threshold of 0.75, consistent with established criteria^33,55^. Colored bars indicate neurons exhibiting robust and statistically significant CP values, whereas grey bars represent neurons whose CP either fell below the 2σ significance threshold or failed to exceed the established CP criterion of 0.75.

### Generalized linear model (GLM)

To predict neural activity from behavior, we trained generalized linear models to predict the activity of each neuron k, as a weighted linear combination of male-directed behaviors (male sniffing and attack behaviors) or food-directed behaviors (food sniffing and eating behaviors), as follows:

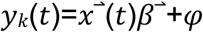

Here, *y*_*k*_(*t*) is the calcium activity of neuron k at time t, 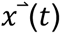) is a feature vector of 3 binary male behaviors at time lags ranging from t-D to t where D = 10s. 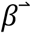 is a behavior filter that describes how a neuron integrates stimulus over a 10s period. *φ* is an error term. The model was fit using 10-fold cross-validation with ridge regularization, and model performance is reported as cross-validated *R*^2^ (cv*R*^2^). To account for cell-cell interactions within the network, we also used the activity of simultaneously imaged neurons as regressors in addition to behavior, as previously described^59^.

### Dynamical system models of neural data

rSLDS models fit the neural data as previously described^46^. In brief, rSLDS is a generative state-space model that decomposes nonlinear time-varying data into discrete states, each with simple linear dynamics. The model describes three sets of variables: discrete states (z) and latent factors (x) that represent the low-dimensional features from the population activity of recorded neurons (y). While the model can also allow for the incorporation of inputs based on behavior features, such external inputs were not included in the first analysis.

The model is formulated as follows: At each time point, there is a discrete state *z*_*t*_ ∈ [1,…, *K*] that depends recurrently on the continuous latent factors (x) as follows:

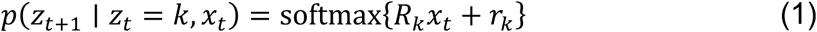

where *R*_*k*_ ∈ ℝ^*K*×*K*^ and *r*_*k*_ ∈ ℝ^*K*^ parameterizes a map from the previous discrete state and continuous state to a distribution over the following discrete states using a softmax link function. The discrete state*z*_*t*_determines the linear dynamical system used to generate the latent factors at any time t:

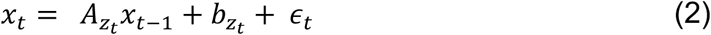

where *A*_*k*_ ∈ ℝ^*d*×*d*^ is a dynamics matrix and *b*_*k*_ ∈ ℝ^*D*^ is a bias vector, where *D* is the dimensionality of the latent space and 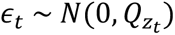 is a Gaussian-distributed noise (aka innovation) term where 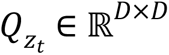.

Lastly, we can recover the activity of recorded neurons by modeling activity as a linear, noisy Gaussian observation*y*_*t*_ ∈ ℝ^*N*^ where *N* is the number of recorded neurons:

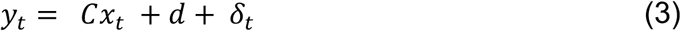

For *C* ∈ ℝ^*N*×*D*^and *δ*_*t*_ ∼ *N*(0, *S*), a Gaussian noise term. Overall, the system parameters that rSLDS needs to learn consist of the state transition dynamics, the library of linear dynamical system matrices, and the neuron-specific emission parameters, which we write as:

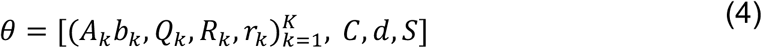

We evaluate model performance using both the evidence lower bound (ELBO) and the forward simulation accuracy (FSA) previously described^46^ and by calculating the variance explained by the model on the data. Briefly, given observed neural activity in the reduced neural state space at time t, we predict the trajectory of population activity over an ensuing short time interval Δt using the fit rSLDS model, then compute the mean squared error (MSE) between that trajectory and the observed data at time t+ Δt. This MSE is computed across all dimensions of the reduced latent space and repeated across cross-validation folds. This error metric is normalized to a 0-1 range in each animal across the holding to obtain a bounded measure of model performance. The FSA can intuit where model performance drops during the recording. In addition to MSE, we also calculate the Pearson’s correlation coefficient (*R*^2^) between the predicted and observed data for each dimension following the forward simulation. By taking the average correlation coefficient across dimensions, we can obtain a quantitative estimate of variance explained by rSLDS on observed data.

The number of states and dimensions used for the model is determined using 5-fold cross-validation. Visualization of the dynamical system using principal components analysis is performed as described previously^46^.

Code used to fit rSLDS on neural data is available in the SSM package: (https://github.com/lindermanlab/ssm)

### Estimation of time constants

We estimated the time constant of each dimension of linear dynamical systems using eigenvalues *λ*_*a*_ of the dynamics matrix of that system, derived previously^40^ as:

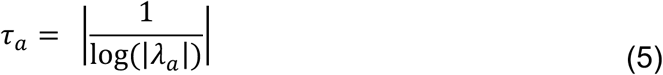

### Calculation of line attractor stability

As a quantitative measure to estimate the variance of approximate line-attractor dynamics, we calculated a log 2 ratio between time constants from X_1_ and X_2_ dimensions as previously defined (sometimes referred to as the “line attractor score”)^46-48,59^ as:

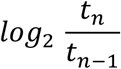

where *t*_*n*_ and *t*_*n*−1_ are the first and second largest time constants of the dynamics matrix.

### Correlation of time-to-peak of integration dimension and latency to first attack

We estimated the linear correlation between the time to peak of the integration dimension or the time to initiate the first attack behavioral action from the time of initiation of sniffing the intruder.

### Calculation of auto-correlation half-width

We computed autocorrelation half-widths by calculating the autocorrelation function for each neuron’s time series data (y_t_) for a set of lags as described previously^22^. Briefly, for a time series (y_t_), the autocorrelation for lag k is:

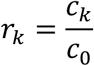

where *c*_*k*_ is defined as:

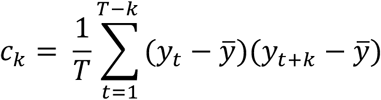

and *c*_0_ is the sample variance of the data. The half-width is found for each neuron as the point where the autocorrelation function reaches a value of 0.5.

### Decoding of behaviors from the rSLDS dimensions

To evaluate whether the integration dimension (X_1_) or X_3_ dimensions could discriminate between attack and eating behaviors, we trained a frame-wise decoder as previously described ^46,55^. Briefly, behavioral "trials" were constructed by concatenating behavioral bouts separated by less than five seconds. We then trained a linear support vector machine (SVM) classifier to determine an optimal decoding threshold for distinguishing the normalized integration dimension signal^46^ during frames associated with attack versus eating behaviors. For shuffled control data, behavioral annotations were randomized across bouts within each trial to disrupt genuine correlations, with this shuffling procedure repeated 20 times per intruder and per mouse. Decoder performance was quantified using the F1 score, averaged across held out "trials" for each animal. Additionally, to rigorously control for potential artifacts arising from slow decay dynamics in the integration dimension, we implemented a permutation test inspired by previous studies^46^. Specifically, a decoding threshold obtained from one mouse was applied to the integration dimension data of all other mice. High performance under this cross-animal decoding paradigm would indicate that decoding accuracy reflects robust behavioral discriminability rather than spurious correlations due to slow drifts in neural activity.

### Quantification and statistical analysis

All statistical analyses were performed with GraphPad Prism 9.0.2. We conducted unpaired two-tailed t-tests to compare two groups that present the normal distribution. In the experiments with paired samples, we used the Wilcoxon matched-pairs signed test or paired t-test. In the experiments with non-paired samples, we used the Mann–Whitney *U* test. For multiple comparisons between the groups representing the normal distribution, one-way ANOVAs followed by Holm Sidak’s post hoc test were performed. The data are presented as the mean ± SEM. One-way or two-way ANOVA tests were followed by Holm Sidak’s post hoc test. The significance threshold was held at a = 0.05, two-tailed (not significant (ns), p>0.05; *p<0.05; **p<0.01; ***p<0.001; ****p<0.0001.

### Reporting summary

Further information on research design is available in the Nature Research Reporting Summary.

## Data Availability

The data used in this study are available at the following URLs: (TBN on acceptance).

## Code Availability

Code for analysis can be found at https://github.com/lindermanlab/ssm. The supporting analysis code will be made available in DANDI upon publication.

---

**Extended Data Fig. 1.**
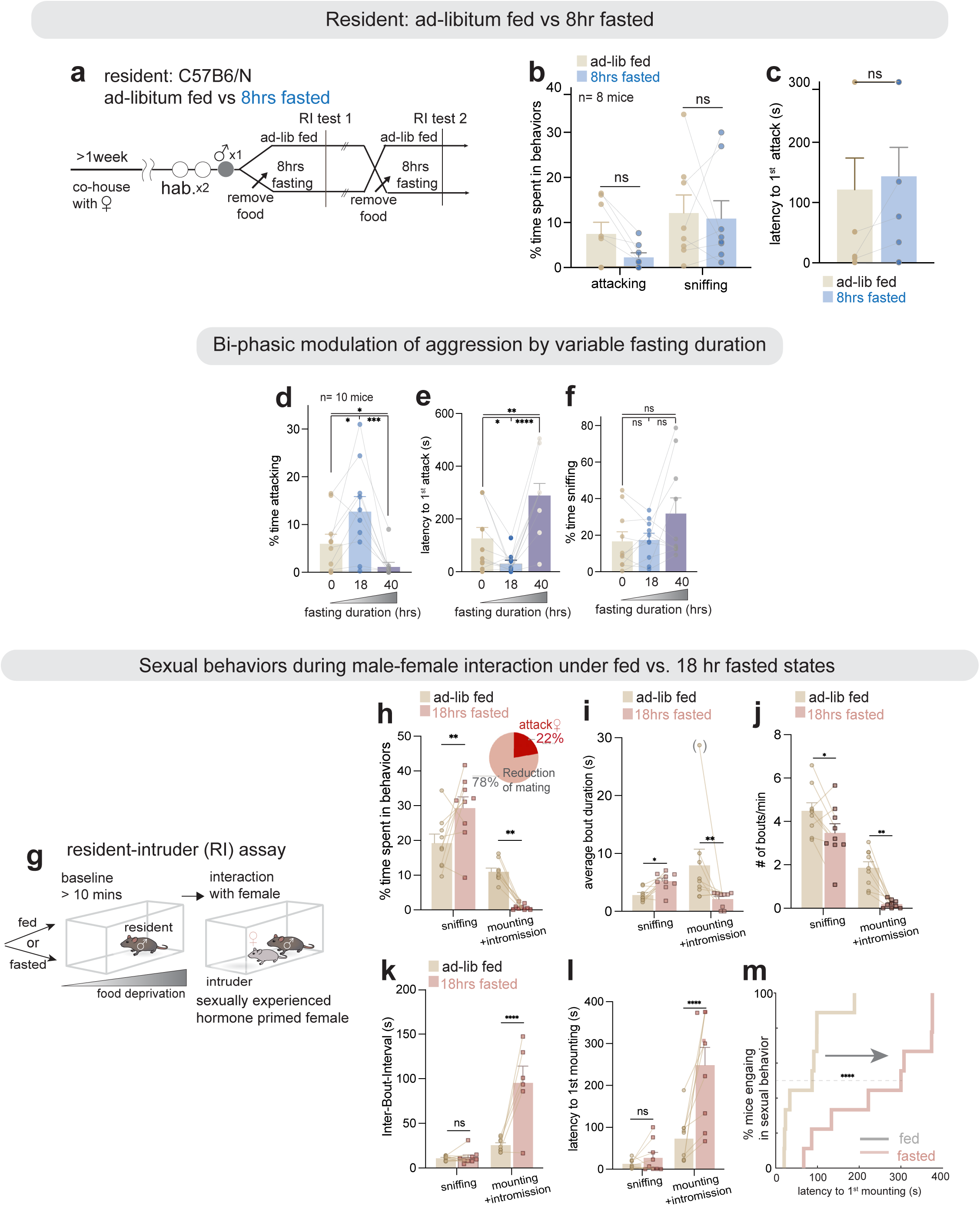
Hunger states modulate innate social behaviors during male-male or male-female interactions; additional information for Fig. 1. (**a-c)** (**a**) Schematic and quantifications of behaviors in fed vs. 8-hour fasted male residents. (**b**) Time spent in attack or sniffing the male intruder (%), and (**c**) latency to the first attack. These males underwent the same conditions as described in Fig. 1, except for 8 hours of food deprivation instead of overnight fasting, with food removal beginning at 9 AM (see Methods for more information, n = 8 mice). (**d-f)** Quantification of aggression parameters across *ad libitum*-fed, 18-hr or 40-hr fasted states: (**d**) time spent in attack (%); (**e**) latency to the first attack from the first sniff; (**f**) time spent in sniffing (%). (n = 10 mice). (**g-m)** (**g**) Sexual behaviors of resident males during interactions with female intruders in *ad libitum*-fed vs 18-hr fasted states; (**h**) quantifications of time spent sniffing females and female-directed mounting or intromission (%). Inset: 22% of males in the fasted state attacked female intruders; (**i**) average bout duration; (**j**) number of bouts per minute; (**k**) Inter-bout interval; (**l**) latency to the first female-directed mounting, and (**m**) distribution showing the percentage of mice engaging in sexual behaviors for the latency to the first female-directed mounting in fed vs. fasted conditions. For (i), one outlier data point with parentheses, since the difference remained statistically significant even with its exclusion. \**p*<0.05, \*\**p*<0.01, \*\*\**p*<0.001, \*\*\*\**p*< 0.0001; ns, not significant. For all figure panels, data are mean ± SEM.

**Extended Data Fig. 2.**
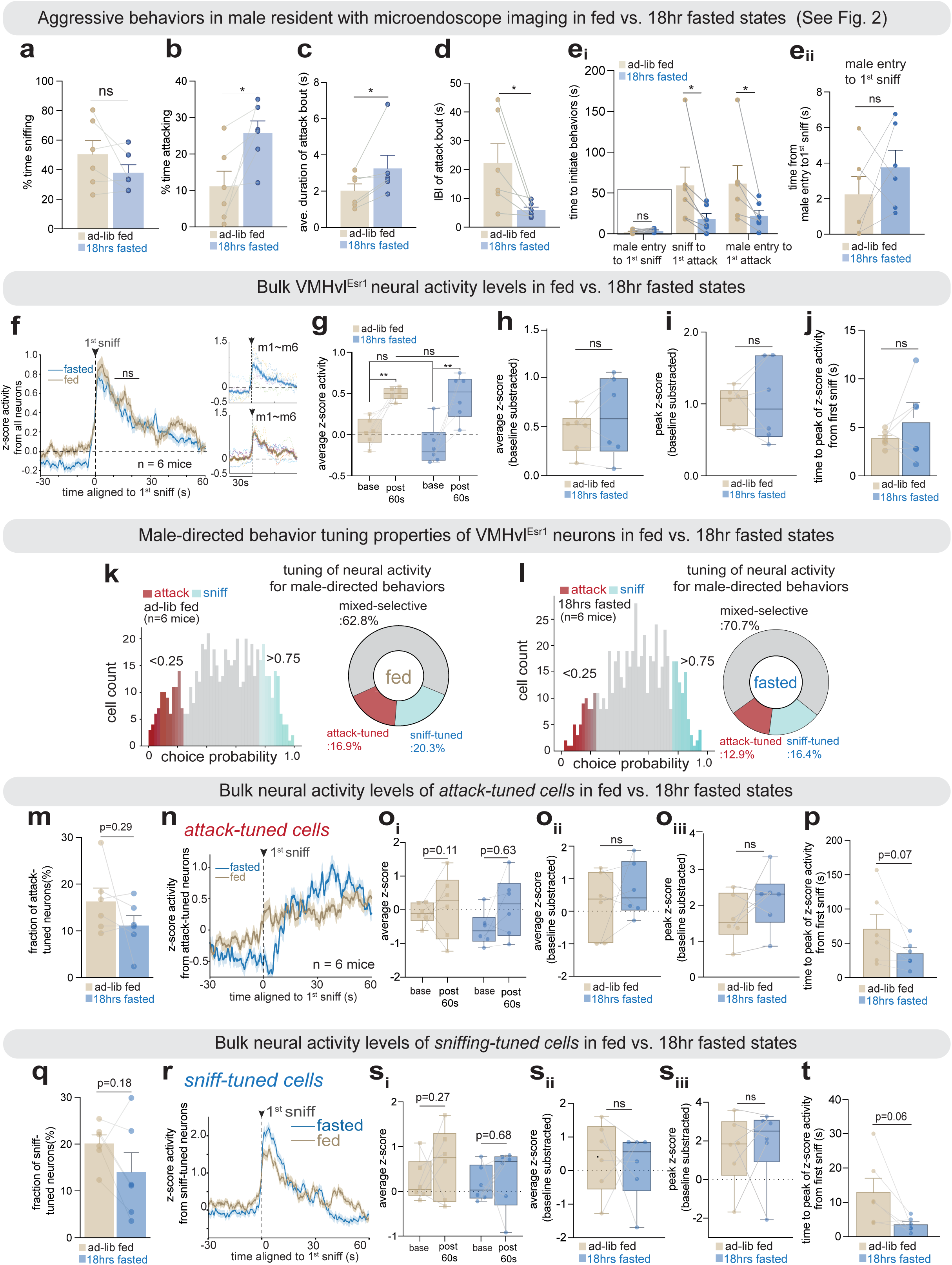
Aggressive behaviors and tuning properties of VMHvl^Esr1^ neurons in *ad-libitum* fed vs. 18-hr fasted states; additional information for Fig. 2. (**a-e)** Quantification of aggression parameters for mice shown in Fig. 2: (**a**) time spent sniffing the male intruder (%); (**b**) time spent in attack (%); (**c**) average attack bout duration; (**d**) Inter-bout interval of attack; (**e**) time from male entry to first sniff and from first sniff to the first attack. **e_ii_**: an expanded panel for **e_i_**-box. (n = 6 mice). (f) (left) BTA of bulk VMHvl^Esr1^ activity aligned to the onset of first sniffing showing no significant difference between *ad-libitum* fed and 18-hr fasted states (n = 6 mice). (right) Dashed lines for individual mice. (**g-h**) (**g**) Average neuronal activity for baseline and 1 min following the first sniff; (**h**) Average baseline subtracted z-score activity between fed vs. 18-hr fasted conditions (ns, p > 0.05). (i) Time to peak z-score following first sniffing (ns, p > 0.05, paired t-test). (j) BTA of bulk VMHvl^Esr1^ activity aligned to the onset of the first attack for between fed vs. fasted states (ns, p > 0.05). (**k-l**) Choice probability for male-directed sniffing vs. attack behavior and percentage of cells with tuning preference in (**k**) fed and (**l**) fasted states. (**m, q**) Fraction of neurons for attack-tuned (left) and sniff-tuned (right) across fed (**m**) and fasted (**q**) states. (**n, r**) BTA of VMHvl^Esr1^ neuronal population tuned to attack vs. sniffing from the choice probability analysis, aligned to the onset of first sniffing between *ad-libitum* fed and 18-hr fasted states (n = 6 mice). *\*p*<0.05; ns, not significant. For all figure panels, data are mean ± SEM.

**Extended Data Fig. 3.**
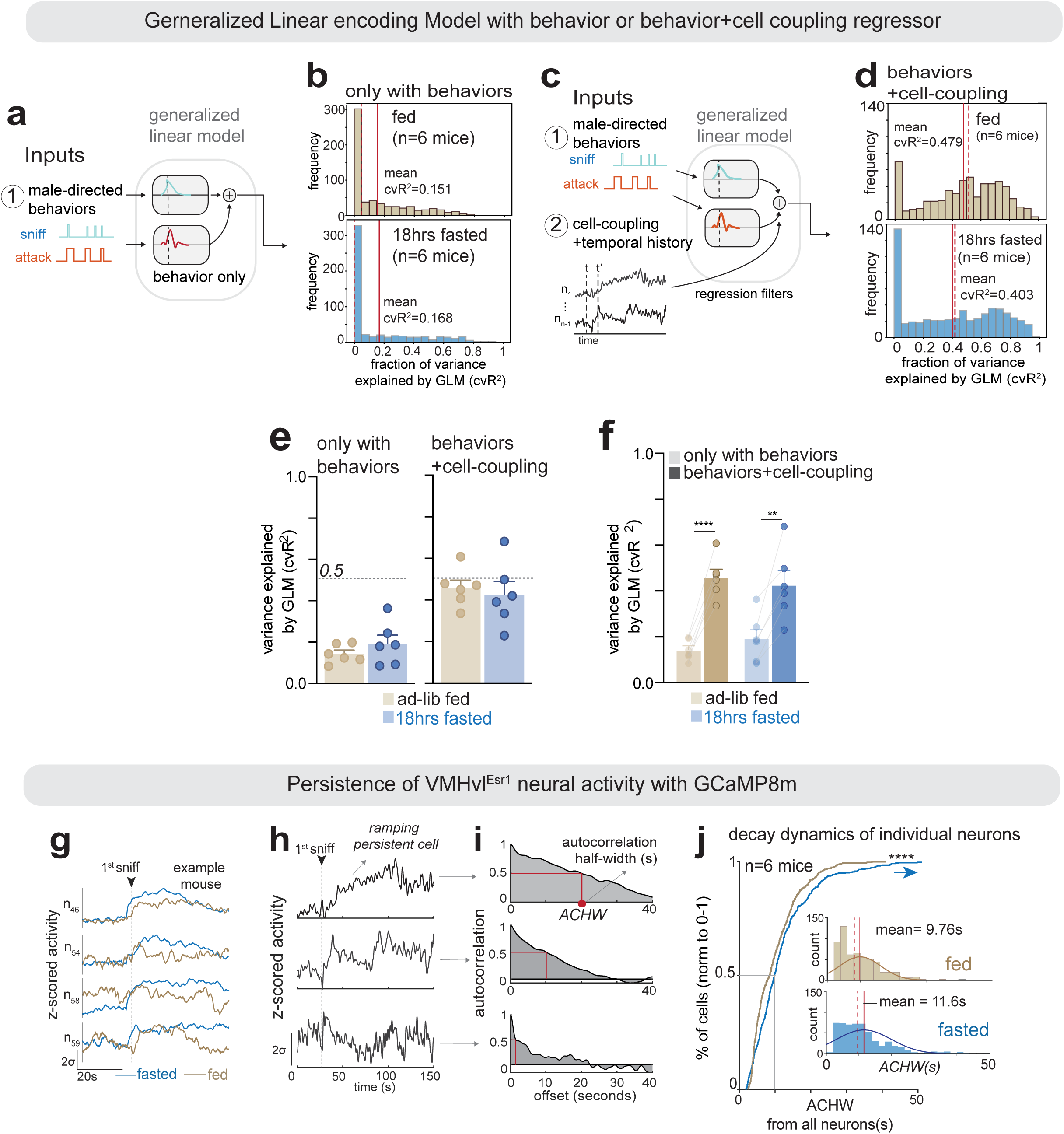
Improved GLM fits with cell coupling and enhanced persistence of VMHvl^Esr1^ neurons; additional information for Fig. 2. (**a**) Schematics of a GLM to predict individual cell activity from a behavior regressor. (**b**) Histogram with variance explained (cvR^2^) across all neurons, n=6 mice from behavior-only GLM in the fed (above) and fasted (below) states. Solid line: mean; dashed line: median value. (**c**) Schematics of GLM to predict individual cell activity from behavior and cell-cell coupling. (**d**) Histogram with cvR^2^ across all neurons, n=6 mice from behavior + cell-coupling GLM in fed (above) and fasted (below) states. Solid line: mean; dashed line: median value. (**e-f**) Quantification of cvR^2^ in GLM with behavior + cell-coupling compared to the behavior-only regressor. (**g**) Individual neural traces between *ad-libitum* fed vs. fasted states, showing persistence during male-male interaction with the GCaMP8m indicator, blue: fasted, brown: fed state. (**h-i**) Individual example neural traces with the most extended (upper, 20s), medium (middle, 10s), and shorter (lower, 5s) ACHW. (**j**) Distribution of cells for ACHW in fed vs. 18-hr fasted state for 6 mice. Histogram of ACHW (inset). Paired or unpaired t-test, *\*\*p*<0.01*, ***p*<0.001, \*\*\*\**p* <0.0001; ns, not significant. For all figure panels, data are mean ± SEM.

**Extended Data Fig. 4.**
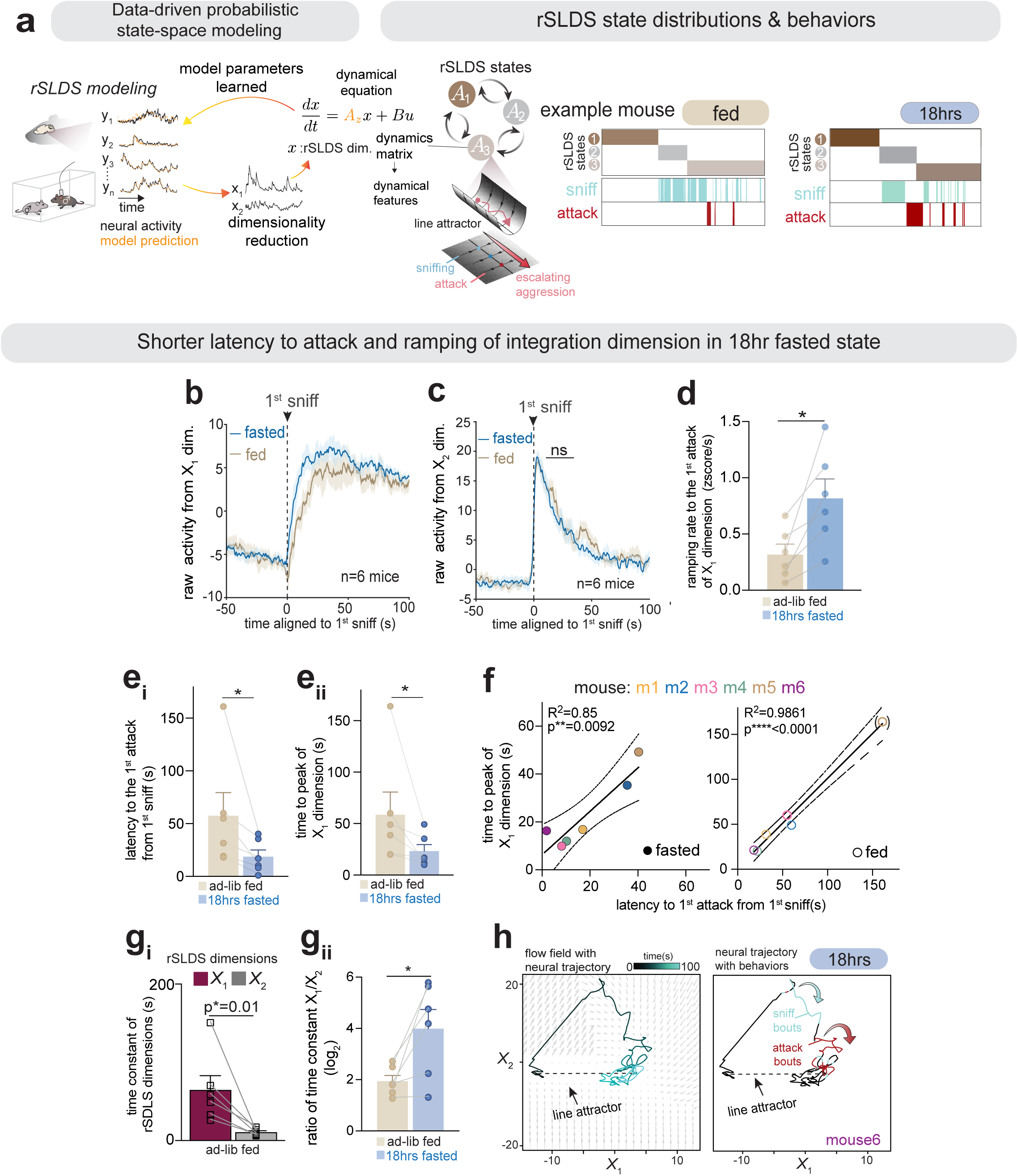
Enhanced ramping and stability of aggression-encoding VMHvl^Esr1^ attractor dynamics in 18-hr fasted state; additional information for Fig. 2. (**a**) (left) Schematics of rSLDS modeling. (right) Discrete states of rSLDS model and behavior raster in Fig. 2f, g from fed and fasted states. (**b-c**) BTA of non-normalized activity in the X_1_ dimension (**b**) or X_2_ dimension (**c**), aligned to the first attack in the fasted and fed conditions (n = 6 mice). Compare with Fig. 2h, i. (**d**) Ramping rate of the X_1_ dimension in Fig. 2jii. (**e**) Quantifications of (**e_i_**) latency to the first attack and (**e_ii_**) time to peak activity in the integration dimension from the first sniff in fed vs. fasted states. (**f**) Correlation between the time to peak integration dimension and the latency to the first attack in the fasted state (left, closed circles) and the fed state (right, closed circles). Individual mice are color-coded. R² = 0.85, **p = 0.0092 (left); R² = 0.98, ****p < 0.0001 (right). One data point with () in the right panel, as the statistical significance did not change even with its exclusion. (**g**) Quantifications of the ratio between time constants from X_1_ and X_2_ dimensions between fed vs. 18-hr fasted. (**h**) Another example of flow field of rSLDS model with neural trajectory in the fasted state. \**p*<0.05, \*\**p*<0.01, \*\*\**p*<0.001; ns, not significant. For all figure panels, data are mean ± SEM.

**Extended Data Fig. 5.**
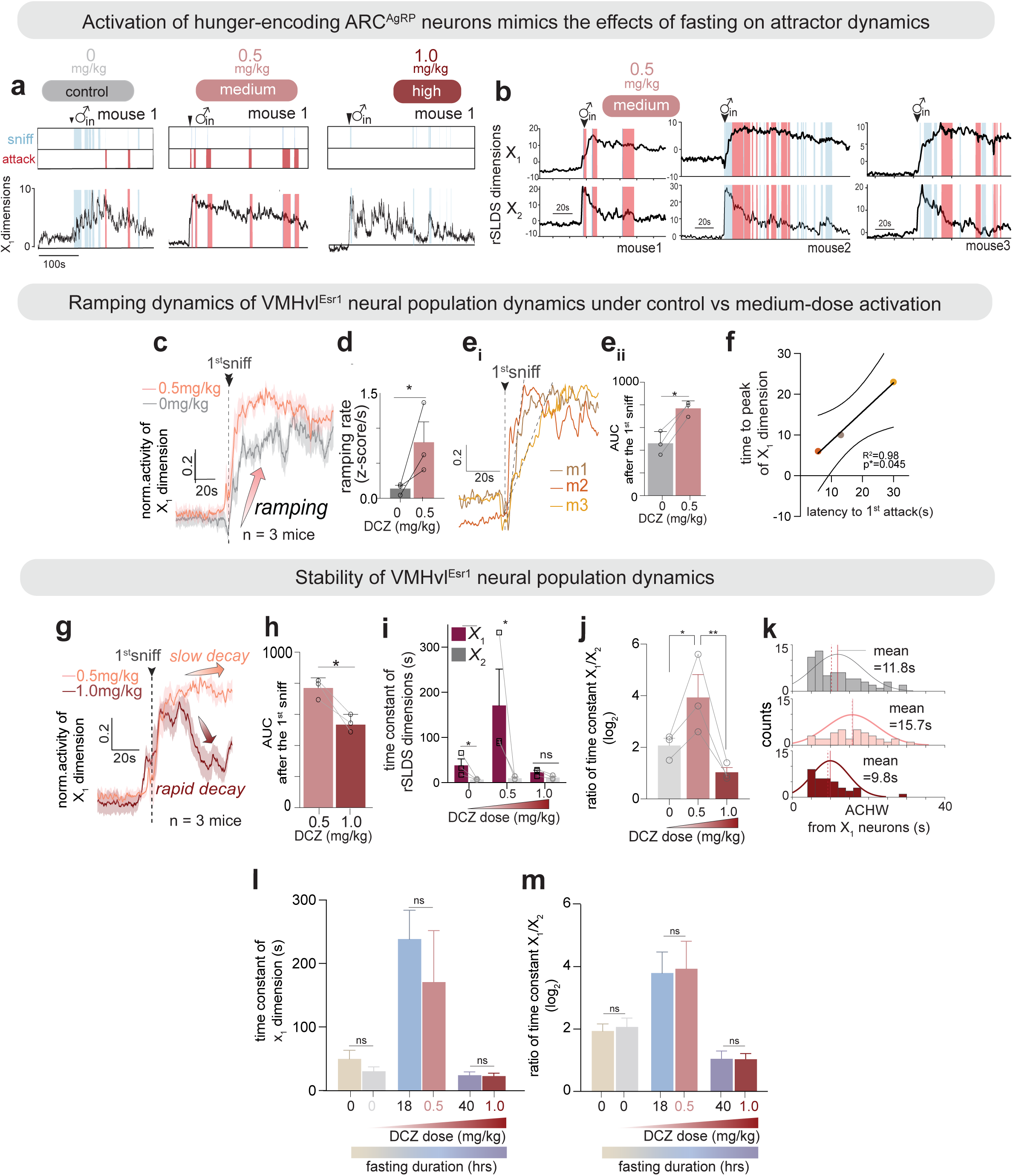
Acute, titrated activation of ARC^AgRP^ neurons shifts aggression-encoding VMH^Esr1^ attractor dynamics; additional information for Fig. 4. (**a**) Representative traces of *X_1_* dimension at each dose for ARC^AgRP^ neuronal activation from a representative mouse with male-directed behavior raster. (**b**) Representative traces of *X_1_* and *X_2_* dimensions in medium-dose conditions showing rapid ramping and persistent activity in the X_1_ dimension with a color-coded raster of sniffing (blue) or attack (red) behaviors across mice. (**c**) BTA of X_1_ dimensions comparing saline (gray) vs medium-dose (pink) conditions, highlighting the accelerated ramping in the medium dose. (**d**) Quantification of the ramping rate from (**c**). (**e**) Individual traces of X_1_ dimension at the medium level of activation with rapid ramping (**e_i_**) and AUC calculated for 100s after the first sniff (**e_ii_**). (**f**) Correlation between the time to the peak of the X_1_ dimension and the latency to the first attack from the first sniff bout. (**g-h**) BTA of X_1_ dimensions comparing medium vs. high-dose condition: (**g**) difference in persistence of the first dimension of rSLDS; (**h**) quantifications of AUC after the first sniff. (**i**) Time constant from X_1_ and X_2_ dimensions across three different conditions of chemogenetic activation of ARC^AgRP^ neurons. (**j**) Quantification of the ratio between the time constants of X_1_ and X_2_ dimensions. (**k**) Histogram of ACHWs across control, medium, and high-dose conditions in Fig. 4j. (**l-m**) Quantification of (**l**) time constant for the X_1_ dimension or (**m**) ratio between time constants for X_1_ and X_2_ dimensions across naturalistic food deprivation or activation of ARC^AgRP^ neuronal activity to induce hunger drive. \**p*<0.05, \*\**p*<0.01; ns, not significant. For all figure panels, data are mean ± SEM.

**Extended Data Fig. 6.**
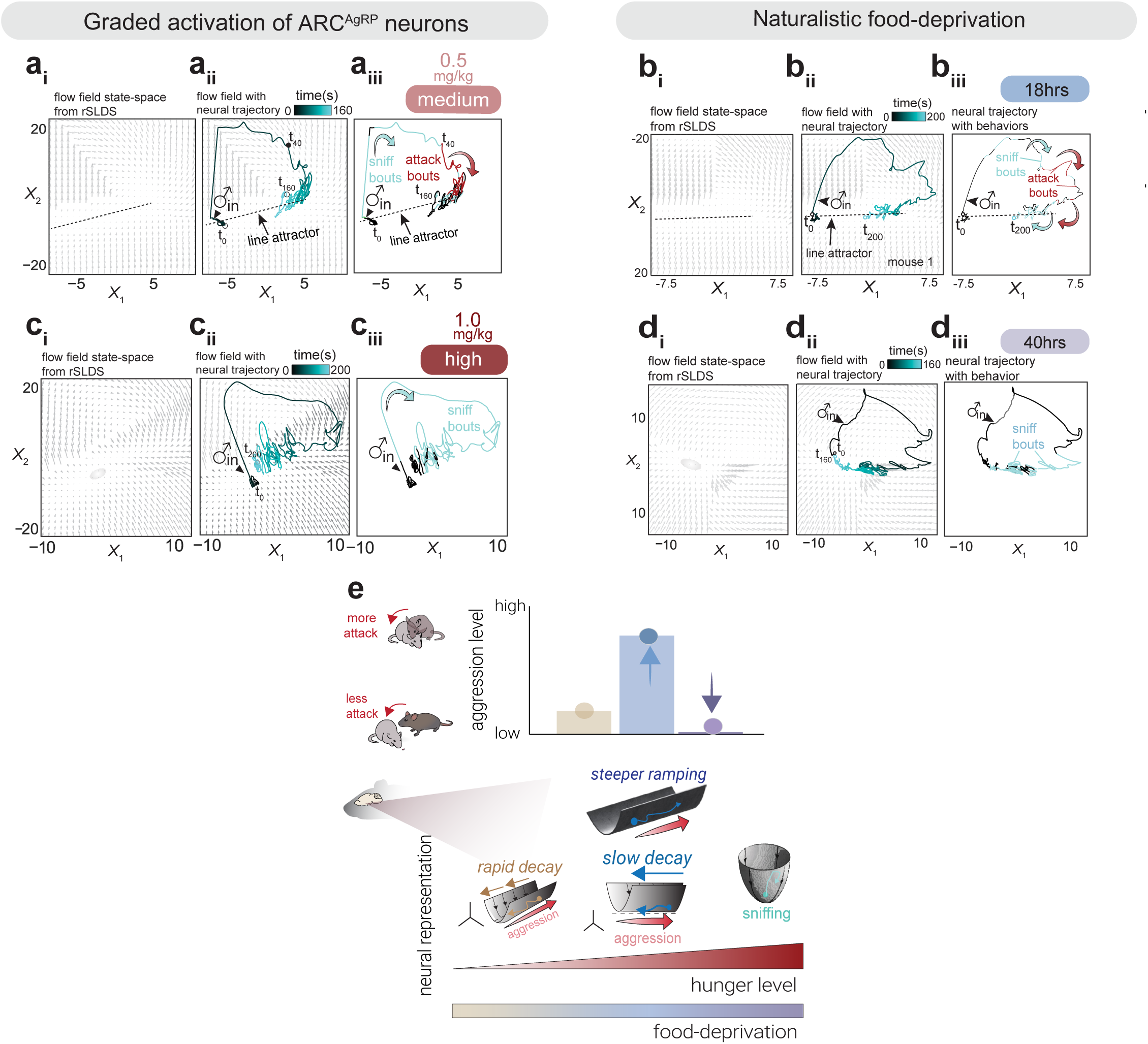
Acute, titrated activation of ARC^AgRP^ neurons phenocopies biphasic starvation-induced shifts in aggression-encoding VMH^Esr1^ attractor dynamics; additional information for Fig. 4. (**a-d**) Representative flow fields with trajectory colored by time (middle) or behaviors (right) from rSLDS-modeled attractor dynamics under naturalistic food deprivation conditions for 18-hr (**a**) or 40-hr (**c**) and chemogenetic activation of ARC^AgRP^ neuronal activity at medium (**b**) or high-dose conditions (**d**). (**e**) Schematic summarizing the effects of various hunger levels on aggression and aggressiveness-encoding VMHvl^Esr1^ neural population dynamics.

**Extended Data Fig. 7.**
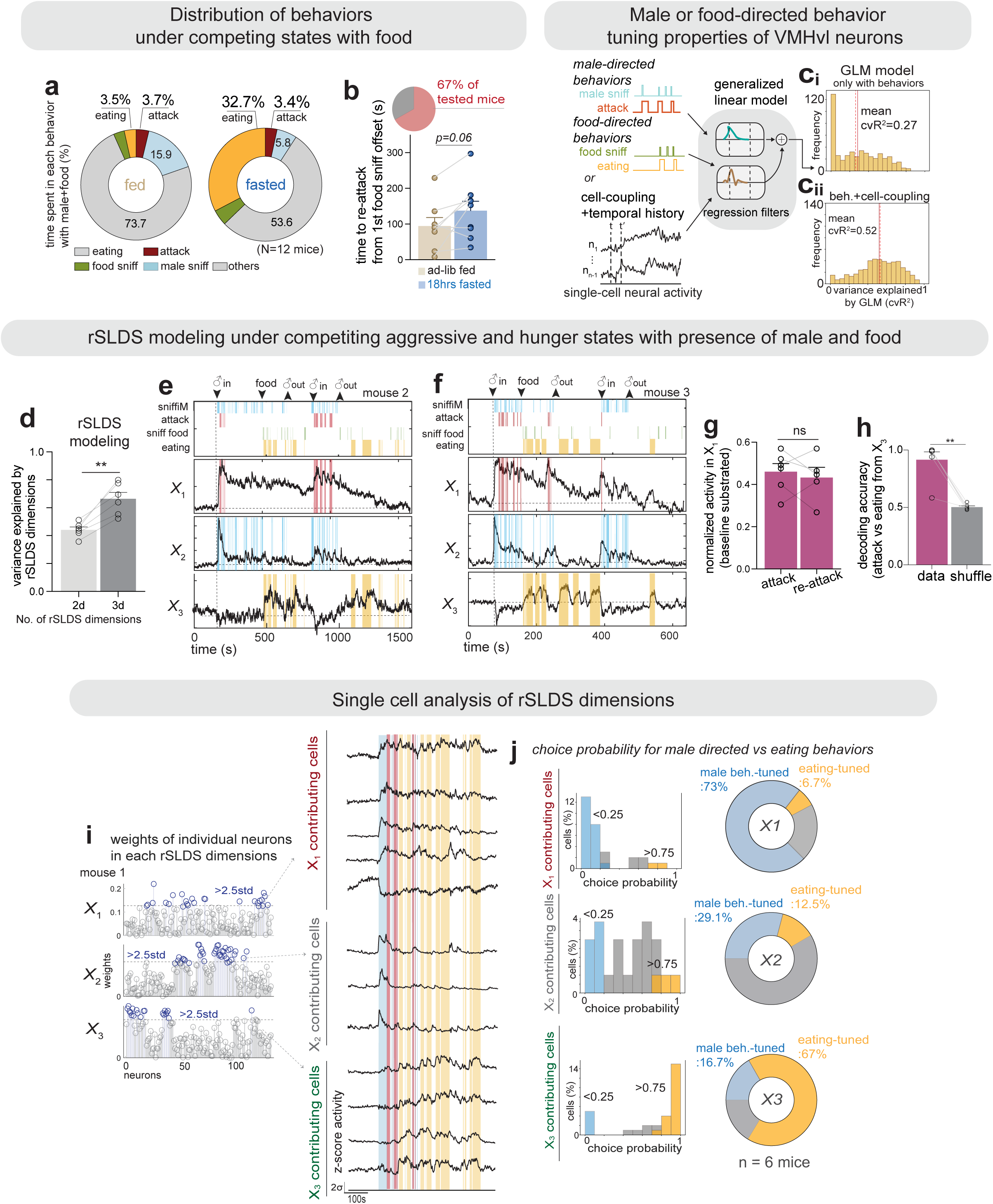
A competing action for food elicits orthogonal perturbations in the VMHvl^Esr1^ attractor dynamics, with distinct tuning properties; additional information for Fig. 5. (**a**) Pie charts of distribution of behaviors when males and food are present together in the fed state (left) and fasted state (right). (**b**) Quantification of time to re-attack from the first food sniff offset in the fasted state. 67% of tested mice exhibited re-attack after transitioning to eating behavior (right). (**c**) Histogram of GLMs with male or food-related behaviors only (**c_i_**) and behaviors with cell-to-cell coupling from the neural activity (**c_ii_**) of the fasting condition. Solid lines: mean values; dashed lines: median values. (**d**) Quantification of rSLDS model performance using two vs. three dimensions. (**e-f**) Representative traces of rSLDS dimensions (upper, X_1_; middle, X_2_; below, X_3_) for (**e**) another mouse and (**f**) the mouse shown in Fig. 5 **f, g**. (**g**) (left) X1-X3 weights for each individual cell in the example mouse used in Fig. 5j. (right) Representative traces of single-cell activity from X_1_, X_2_, and X_3_-weighted neurons. (**h**) Choice-probability of male-directed behaviors (attack + sniff) vs. feeding (eating) in the fasted state in X_1_, X_2_, X_3_-weighted neurons (n = 6 mice). (**i**) Quantification of X_1_ activity after attack or re-attack, reused from Fig. 5mi and 5mi**_v_**. (**j**) Decoding accuracy distinguishing eating vs. attack bouts in real vs. shuffled data from the X_3_ dimension (n = 6 mice). \*\**p*<0.01; ns, not significant. For all figure panels, data are mean ± SEM.

**Extended Data Fig. 8.**
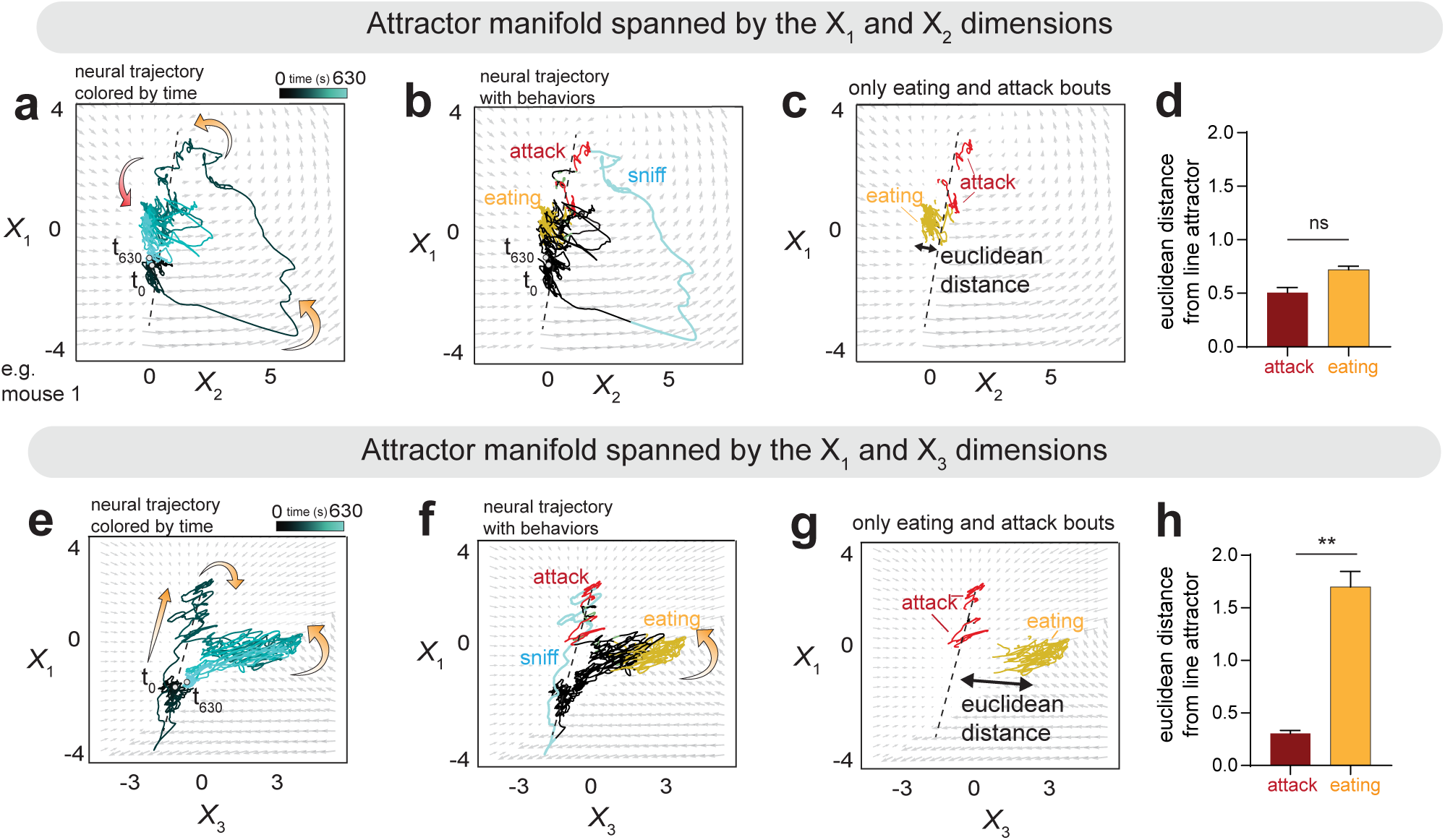
Trajectories in rSLDS state space reveal distinct manifolds, with a persistent and attractive aggressive state under drive for feeding in the VMHvl^Esr1^ neural population; additional information for Fig. 5. (**a-c**) rSLDS modeling based on the state space of the X_1_-X_2_ dimensions with trajectories (**a**) colored by time, (**b**) by behavior, and (**c**) trajectories during attack and eating behavior bouts. (**d**) Quantification of Euclidean distance in Fig. 8c from the line attractor for attack and eating behavior bouts. (**e-g**) Neural state space of the X_1_-X_3_ dimensions with trajectories: (**e**) colored by time, (**f**) colored by behavior; (**g**) only trajectories during attack and eating. (**h**) Quantification of Euclidean distance in Fig. 8g from the line attractor during attack and eating behavior bouts. \*\**p*<0.01; ns, not significant. For all figure panels, data are mean ± SEM.

## Notes

### Competing Interest Statement

The authors have declared no competing interest.

### Summary of Updates

Corrected the typo in one of the authors' names as it appeared in the online system.

